# Multi-platform evaluation and optimization of single-cell RNA isoform sequencing

**DOI:** 10.1101/2025.02.22.639662

**Authors:** James T. Webber, Daniel A. Bartlett, Ghamdan Al-Eryani, Akanksha Khorgade, Asa Shin, Christophe Georgescu, Houlin Yu, Brian J. Haas, Victoria Popic, Aziz Al’Khafaji

**Affiliations:** Broad Clinical Labs, Cambridge, MA; Broad Institute of MIT and Harvard, Cambridge, MA

## Abstract

Long-read transcriptomics enables isoform identification and quantification at single-cell resolution, but analysis is complicated by artifacts introduced during library preparation. We characterize the distinct artifact profiles of three popular single-cell platforms and demonstrate their impact on isoform identification. We introduce a new method for the identification of a wide range of artifacts in cDNA libraries and novel biochemical and bioinformatic strategies to significantly reduce their impact on downstream applications. Finally, we provide a comprehensive framework for RNA isoform sequencing analysis and interpretation as the field continues to develop.

## Main

Advances in long-read sequencing platforms, library construction methods, and computational tools have collectively turned single-cell isoform sequencing into a production-level assay for biological research. The identification and quantification of RNA isoforms at the single-cell level allows researchers to resolve alternative splicing across cell types. Alternative splicing is a central regulatory axis which modulates gene function through differential exon splicing during transcript maturation. In contrast to gene expression analyses, full-length transcript expression analyses allow researchers to capture the rich structural information embedded in the length of the transcript, enabling differentiation of active and inactive forms of the same gene, identification of localization signals, and discovery of novel isoforms. While this datatype is feature rich, artifacts that arise during library construction and sequencing can be erroneously assigned to novel transcripts, greatly complicating analysis. In this work, we identify key sources of artifacts in single-cell RNA isoform sequencing data and implement molecular and computational strategies to eliminate or minimize their impact on downstream analysis.

Biochemical and workflow differences between single-cell platforms can lead to important differences during the generation of the sequencing libraries, and thus the generated data. To enable accurate isoform-level transcriptomics it is critical to characterize and account for the landscape of artifacts produced from various single-cell approaches. Artifacts arise during the processes of 1) cDNA synthesis and PCR amplification due to off-target priming from reverse transcription (RT), template switch oligo (TSO), and PCR primers; 2) internal priming from oligo(dT) RT primers during reverse transcription, leading to unintended priming at A-rich regions; 3) mRNA degradation during library construction; 4) incomplete reverse transcription; and 5) errors in sequencer primary data processing^1,2^. These artifactual reads may be difficult or impossible to confidently assign to the originating isoform, which can lead to inaccuracies in quantification and the erroneous assignment of novel isoforms.

We evaluated multiple prominently used single-cell platforms, namely PIPseq V4 (Fluent Biosciences), 10x 3′ V3.1 (10x Genomics), and 10x 5′ V2.1 (10x Genomics), for their ability to generate high-quality full-length cDNA libraries for isoform sequencing^3^. Given the biochemical differences between these three platforms, we hypothesized that they would exhibit notable differences in the transcripts captured and artifacts produced, which can ultimately lead to significant performance variation in downstream isoform identification and quantification. We first prepared single-cell cDNA libraries from peripheral blood mononuclear cells (PBMCs) and sequenced them using short-read sequencing (Illumina Novaseq X). After subsampling to an equal number of reads per cell and clustering each dataset, we found that the three technologies mostly captured the same cell types at similar proportions (Fig. 1a, b, Supplemental Table 1), with two notable differences. Analysis of the 10x 3′ data resolved a population of dendritic cells that were not discretely identified by the other two platforms, while the 10x 5′ and PIPseq V4 platforms were able to resolve innate lymphoid cells. This variation is likely due to differential capture of transcript biomarkers. In addition, the PIPseq library contained a substantially smaller proportion of CD14+ monocytes (8.4% of cells, compared to ∼22% in 10x 3′ and 5′), and the UMIs/cell for CD14+ monocytes was lower than for other cell types. This is consistent with previous reports of this platform, and reflects increased RNAse activity due to cell stress during sample preparation^4^. Overall, the 10x libraries captured more UMIs/cell (median 10x 3′: 10,091; 10x 5′: 6,804; PIPseq: 3,872) and genes/cell (median 10x 3′: 2,904; 10x 5′: 2,339; PIPseq: 1,398), while PIPseq captured more cells overall (10x 3′: 3,687; 10x 5′: 3,083; PIPseq: 8,079; Supplementary Fig. 1). We performed a barnyard experiment using the PIPseq platform and observed that the ability to capture more cells did not come at the cost of higher doublet rates (0.9% of 17,050 barcodes observed with ≥10,000 UMIs; Supplementary Fig. 2), and the ambient contamination was similar across all three methods (median 1% cross-species UMI contamination; Supplementary Fig. 3).

**Figure 1.**
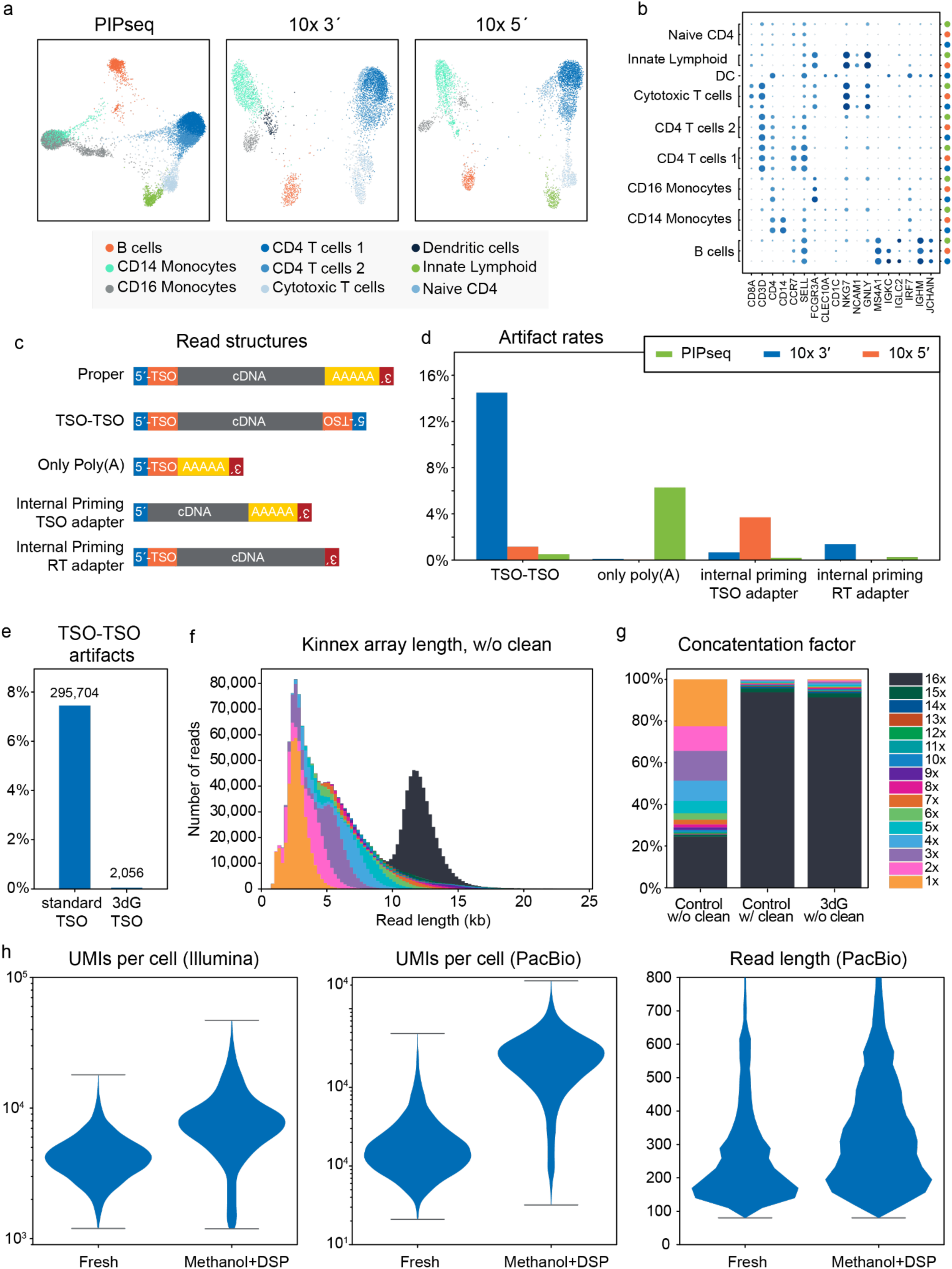
Cross-platform artifact quantification and evaluation of molecular optimizations to minimize their formation. **a)** Single-cell embeddings for PIPseq, 10x 3′ and 10x 5′ libraries prepared from a sample of PBMCs. Cells are colored based on cluster assignment. **b)** A dot plot showing the percentage non-zero (size) and mean expression (color intensity) of a set of markers across clusters. **c)** Schematic showing several common structural artifact classes that `marti` will search for when annotating long reads. The specific sequences of the adapters are input in a configuration file. **d)** Artifact counts and percentages for the most common categories in the PBMC monomer data. Structural annotations are available in the supplementary data. Samples are colored as in 1b. **e)** rate of TSO-TSO artifacts in a standard 10x 3′ monomer library, compared to the 3dG monomers. **f)** The distribution of read length and array size for a 10x 3′ Kinnex library when artifact clean-up is not performed. **g)** Array size distributions for 10x 3′ Kinnex libraries using three protocols: using a 3dG TSO without cleanup, using the standard 10x TSO without cleanup, and using the standard TSO with the cleanup step. **h)** Comparisons for PIPseq data when using fresh PBMCs vs methanol+DSP fixation of the same sample. Violin plots show the number of UMIs per cell from the short-read and long-read sequencing along with the read length distribution for CD14+ monocytes.

We then turned to comparing cross-platform isoform capture. To evaluate the structure of the cDNA library we sequenced the pre-fragmented full-length cDNA libraries as monomers using the Iso-Seq method on a PacBio Sequel IIe sequencer. We also performed MAS-seq using the commercially available Kinnex single-cell kit, and sequenced on the PacBio Revio platform to obtain a deeply sequenced dataset of single-cell isoforms (cDNA reads after deconcatenation: 10x 3′: 91.8M; 10x 5′: 69.7M; PIPseq: 101.3M, Supplementary Fig. 4, 5). We annotated the monomer reads using `marti`, a novel software framework we developed for the in-depth classification and quantification of artifactual cDNA constructs. `marti` enables the detection of a wide range of artifactual constructs that arise during cDNA synthesis, PCR, sequencing, and primary sequence analysis stages (Fig. 1c, d; Supplementary Table 2; methods). By examining the occurrences of the RT and PCR primers, sequencing adapters, and poly(A) tails within the read sequence, `marti` builds the structural representation of the read and classifies it based on its deviation from the non-artifactual (proper) read conformation and its matches against a library of predefined artifactual constructs. Notably, by reporting the specific structure of each read (in addition to the matched artifact class(es)), `marti` facilitates the detection of novel artifacts not yet defined in its artifact library. We used `marti` to quantify the rate of occurrence of different artifact types in our long-read libraries, and extract the annotated native cDNA sequence of proper reads for alignment and downstream analysis.

Generally, artifacts can fall into two non-exclusive classes, those with an improper adapter structure (e.g. TSO-TSO, RT-RT, internal PCR adapter priming) and those with evidence of oligo(dT) internal priming, which are structurally indistinguishable from proper non-artifactual reads (Supplementary Fig. 6). Both classes of artifacts are problematic as they do not faithfully reflect the native mRNA and constrain our ability to accurately identify and quantify transcripts. While easier to classify, artifacts with improper adapter structure are particularly problematic as they can interfere with library construction and require involved purification. As seen in previous studies, we observed that the 10x 3′ library contained a high percentage (14.50%) of TSO-TSO artifacts, while the 10x 5′ and PIPseq libraries had lower rates (1.16% and 0.51% respectively)^5^. These and other artifact types inhibit the formation of full-length MAS-seq arrays by introducing a segment with symmetric ends, reversing the array ordering and creating shorter arrays (Fig. 1f).

As a strategy to prevent TSO-TSO artifact formation, we compared the use of the standard TSO with one incorporating 3′-Deoxyguanosine (3dG) in the 10x 3′ Gene Expression kit with PBMCs as input. This modified template switching oligo retains its templating ability without having the requisite 3′-hydroxyl group which can erroneously prime RNA and extend during the reverse transcription reactions. After sequencing these libraries on the Sequel IIe and annotating the reads with `marti`, we found that the 3dG reagent successfully prevents the formation of TSO-TSO artifacts: < 0.1% of reads, compared to 7.45% of reads for the standard TSO in this experiment (Fig. 1e).

We found the 3dG oligo to be highly effective for standard 10x 3′ library construction and sequencing, and, in fact, to result in a reduced number of intergenic and antisense reads found in both short-and long-read data (Supplementary Table 3). While the standard TSO yielded more UMIs/cell overall, further inspection of both TSO conditions revealed a similar number of forward-strand exonic reads, suggesting the potential for elevated artifact rates from the standard TSO. Such artifactual reads have the potential to negatively impact isoform identification and quantification. In addition to the observed improvements in data quality, the 3dG library could be used to create full-length MAS-seq arrays without the need for involved cleanup steps, (91.36% of the arrays were full-length, compared to 24.51% for the standard TSO without cleanup and 93.78% with cleanup; Fig. 1g, Supplementary Fig. 7).

The PIPseq library suffered from a large number of reads that contained only a poly(A) tail and no other sequence, and a lower percentage of full-splice match reads, consistent with the mRNA degradation within the CD14 monocytes found in the short-read data (Fig. 1d, Supplementary Fig. 8, Supplementary Table 4, 5). While these reads do not interfere with the construction of MAS-seq arrays, they reflect a limitation of the PIPseq workflow when capturing cell types with high RNAse levels. Previous work has shown that cell fixation prior to PIPseq encapsulation can prevent mRNA degradation, which would preserve more full length molecules in these cells^6,7^. To explore this, we tested five different fixation methods on a new PBMC sample and used both short and long-read sequencing to evaluate the results. We found a combination of methanol and dithio-bis(succinimidyl propionate) (DSP) was most effective at preserving mRNA from CD14+ monocytes. Kinnex single-cell sequencing of the PBMCs showed that methanol DSP fixation improved CD14+ monocytes as measured by the median number of reads per cell (fixed: 2,253; fresh: 153), UMIs per cell (fixed: 1,933; fresh: 135) and the median read length (fixed: 305bp; fresh: 223bp) (Supplementary Fig. 9, 10). While the fixed condition does produce higher quality PIP-seq data, the modest monocyte read lengths are likely due to cellular stress conditions during the involved cell preparation for the single-cell experiment.

While addressing the biochemical root cause of artifacts is the ideal path, such workflow fixes can take extensive development and optimization. Such is the case for internal priming of the oligo(dT) during reverse transcription, wherein a stretch of adenines in a transcript is sufficient for priming and extension. These internal priming events are upstream of the poly(A) tail and hence truncating. During analysis, these reads appear to be novel 3′ sites, either for known genes (leading to mistaken annotation of novel isoforms) or in intergenic regions (leading to mistaken annotation of novel genes). Identifying internally primed reads can be challenging because they appear structurally sound and have all expected adapters. In advance of molecular solutions for internal priming in high-throughput single-cell workflows, we developed a workflow to detect internal priming that classifies reads according to the genomic context of their alignment, similar to previous approaches to this problem ^8,9^. Aligned reads were classified based on the proximity of the 3′ end to known transcription stop sites and other annotated features, as well as the genomic context. To be classified as an expected priming event, a read must terminate near a known transcription stop site or be adjacent to a poly(A) signal motif, and must not be located within a coding sequence or be adjacent to a particularly adenine-rich genomic sequence (see methods, Supplementary Table 6). While these heuristics are imperfect, this approach allowed us to broadly classify reads into good priming (known sites and unannotated) and likely internal priming candidates (genomic poly(A)). We observed a striking degree of internal priming across the three platforms (PIPseq: 18.27%, 10x 3′: 26.76%, 10x 5′: 18.69%; Fig. 2a, Supplementary Table 7) and explored the downstream analytical impacts in the next section.

**Figure 2.**
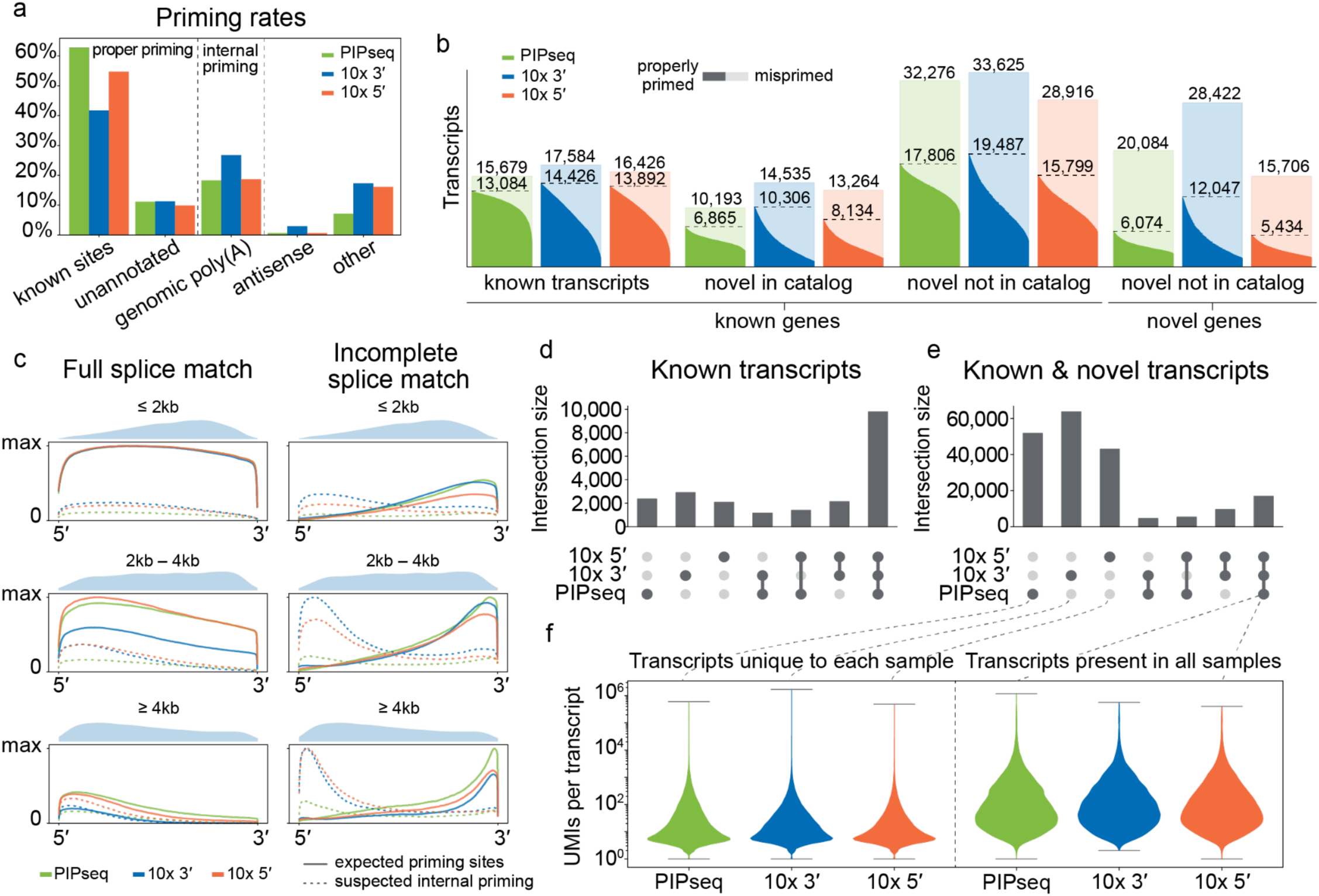
Cross-platform quantification of internal reverse transcription priming artifacts and their impact on downstream analysis. **a)** Rate of different priming events in single-cell MAS-seq data. A complete breakdown of read classifications and rates is available in the Supplementary Table 6. **b)** Visualization of all transcripts identified by `IsoQuant` showing the proportion of reads assigned to that transcript that were classified as properly primed (dark color fill) or misprimed (light color fill). Transcripts are broken down by SQANTI3 class, from left to right: known transcripts present in the reference, NIC transcripts from known genes, NNIC transcripts from known genes, and transcripts assigned to novel genes. Transcripts are displayed horizontally with the dark color fill portion reflecting the proportion of reads for the transcript that came from proper priming sites. **c)** Coverage from full splice match and incomplete splice match reads for known transcripts, binned by transcript length and broken down by priming call. The shaded area above each plot shows the locations for the most-3′ splice junction in each length bin. **d)** Upset plot showing the overlap of known transcripts identified in the three libraries by `IsoQuant`, as determined by `gffcompare` matched as `=` and `c`. **e)** Upset plot showing the overlap of all transcripts. **f)** Violin plot showing the UMIs per transcript for categories in 2e. On the left is shown the UMI distributions for transcripts unique to each sample. On the right, the distributions for transcripts found in all three samples.

After classifying the priming status of each read, we quantified the proportion of reads for each transcript that derived from expected priming sites (Fig. 2b). In all three experiments we saw that the identification of many novel transcripts and novel genes had been derived entirely from internally primed reads (NIC: 29.1% – 38.7%; NNIC: 42.0% – 45.4%; novel genes: 57.6% – 69.8%), raising questions about the reliability of these novel annotations. Known transcripts were more likely to be represented by an expected priming site, but in some cases the only evidence for a known transcript came from internal priming (15.4% – 18.0%). These results highlight the difficulty of accurate transcript identification and quantification in the presence of internally primed reads.

The impact of internal priming can be seen clearly in the average transcript coverage in our MAS-seq experiments. In each experiment we calculated the transcriptome coverage profile by computing the coverage per transcript, normalizing to the length of the transcript, and then averaging the normalized profiles together (Fig. 2c, see methods section). This coverage profile was calculated for four combinations of priming status (good or internally primed reads) and transcript match (full splice match or incomplete splice match). A full splice match (FSM) matches all of the annotated splice junctions for a transcript, while an incomplete splice match (ISM) matches a subset–the read is either 5′ or 3′ truncated, or both. In all three experiments we see that coverage for FSM reads is mostly uniform over the transcript body. As transcript length increases, there is a 5′ bias in FSM coverage, reflecting that a read does not need to contain the entire terminal 3′ exon to be classified as a full splice match, irrespective of internal priming. Long transcripts often contain very long 3′ UTR sequences in the final exon (Fig. 2c, shaded regions), resulting in the observed increased 5′ coverage bias in FSMs compared to shorter transcripts. Internal priming sites in the terminal exon can still produce a full splice match, and we observe this across all three platforms. Finally, reads from a shorter isoform may be erroneously assigned to a longer version. This can be due to unannotated alternative polyadenylation sites in the 3′ UTRs of these genes, or an unexpected combination of transcription start site, exon splicing, and polyadenylation site (Supplementary Fig. 11)^10–12^.

The coverage for incomplete splice matches looks notably different. In all three experiments we see a strong 3′ bias, reflecting reads that were properly primed but failed to capture the whole mRNA. Roughly 50% of the coverage from these properly primed reads fell in the last 25% of the transcript body (10x 3′: 51.8%; 10x 5′: 49.6%; PIPseq: 51.7%; see Supplementary Fig. 12). This can occur because reverse transcription did not produce a full-length cDNA, or because the mRNA itself was truncated. For the longest transcripts (>4kb) the effect is more pronounced (10x 3′: 56.3%; 10x 5′: 54.1%; PIPseq: 52.4%), suggesting that cDNA synthesis is a limiting factor when sequencing these isoforms. On the other end, the most 5′ quartile of longest transcripts accounts for a large proportion of the internally primed ISM coverage (10x 3′: 45.3%; 10x 5′: 45.1%; PIPseq: 32.2%). This reflects that internal priming sites in the middle of a transcript will produce a 3′-truncated read that is missing later splice junctions. This effect is visible in all three experiments but is more pronounced in the 10x 5′ and 3′ data, reflecting the higher rate of internal priming in those libraries.

Our hypothesis was that differences in capture and internal priming behavior between these three platforms would mostly lead to differences in novel isoform calls, while known transcripts would be identified more consistently. To test this we processed each MAS-seq dataset with the isoform identification and quantification tool `IsoQuant` (v3.4.2), and compared the resulting annotations using `gffcompare` (v0.12.6)^13,14^. When considering only isoforms that are closely compatible with known annotations (`=` and `c` classifications from `gffcompare`), there is a strong overlap between all three platforms, suggesting that these isoforms are truly expressed in the sample and that all three platforms capture those transcripts (9,784 transcripts in common; Fig. 2d). However, when we consider all isoforms reported by `IsoQuant`, we observe a very large number of transcript structures that are unique to each of the three single-cell approaches (10x 3′: 63,373; 10x 5′: 43,017; PIPseq: 51,790; Fig. 2e). These transcripts are typically supported by a smaller number of reads (median 10 – 13 UMIs per unique transcript, compared to 67 – 76 UMIs for shared transcripts) and these reads are less likely to be coming from expected priming sites in the genome (27.4% – 37.8% of reads had expected priming, compared to 83.2% – 92.8% for shared transcripts) which supports the hypothesis that misprimed reads can lead to the identification of unreliable novel isoforms (Fig. 2f).

In this study we quantified the rate of common artifacts in single-cell RNA libraries, demonstrated how they can lead to inaccuracies in isoform identification and quantification, and presented strategies to mitigate these issues. In some cases sequencing artifacts can be identified by inspection of the read structure, while other cases must be inferred by considering the genomic context of the aligned sequence. Structural artifacts can be filtered out bioinformatically, and we developed `marti` to identify and remove these artifacts. Preferably, improvements in biochemistry would prevent these artifacts from forming in the first place. We demonstrated the impact of using the 3dG template switch oligo, which prevents a common class of sequencing artifacts in 10x 3′ libraries. We also showed that cell fixation is critical for maintaining mRNA integrity on the PIPseq platform. Other artifacts, such as reads with proper structure arising from internal priming, are more difficult to filter due to our incomplete knowledge of the transcriptome and the aim to discover novel genes and isoforms in our experiments. We developed workflows to tag reads with their priming status and annotate isoforms with their support from both properly and internally primed reads. These annotations can be coupled with custom filters to minimize spurious isoform detection.

As biochemical and bioinformatic methods continue to advance, the best strategy for these data will continue to evolve. Future advances in biochemistry may be able to prevent other classes of sequencing artifacts from being generated, greatly simplifying identification and leading to more accurate and reliable isoform expression results. In particular, more reliable capture at the poly(A) tail would reduce the sequencing of internally primed reads and more processive reverse transcriptases would provide more complete coverage over long transcripts. While methods exist to sequence specifically from the 3′ end of mRNA molecules, they have yet to be demonstrated at the scale and throughput necessary for single-cell isoform sequencing^15^. Advances in throughput and accuracy of full-length RNA isoform sequencing will further bolster a more comprehensive representation of transcriptomics and bring to light critical roles RNA splicing plays in natural and disease biology.

## Online Methods

### Wet-lab

#### PIP-seq vs 10x 3′ vs 10x 5′ Fresh PBMC sample preparation

Cryopreserved PBMCs obtained from AllCells (Lot # 3082436 25/Apr 22 with 10 million cells) were thawed for 2min in 37°C water bath and washed twice using PBS containing 0.4 % BSA, paletted using a swing bucket centrifuge (300 x g for 5 minutes, 4°C). Cells were then resuspended in Fluent cell resuspension buffer (FB0002440), filtered through 40um Flowmi cell strainers (Bel-Art, H13680-0040) counted and viability evaluated with an AO/PI stain (90.5% viability) using Nexcelom Cellometer Auto 2000 (Nexcelom Bioscience). The cell suspensions were mixed well and were divided equally into three aliquots, two for the 10x Chromium captures (3′ and 5′ assays) and one for PIP-seq captures. Samples were further diluted using a Fluent cell resuspension buffer to meet manufacturer requirements for cell loading concentration; 800 cells/ul for both 3′ and 5′ 10x Chromium captures and 1650 cells/ul for PIP-seq capture.

#### Barnyard K562:3T3 sample preparation

K562 (ATCC) and 3T3 cells (ATCC) were harvested and washed twice in PBS containing 0.4% BSA, palleted at 300 x g for 5 minutes in a swing bucket centrifuge (4°C). Cell pellets were resuspended in 400 µl of room-temperature Fluent Cell Resuspension Buffer (FB0002440) and analyzed for cell count and viability using a Nexcelom Cellometer Auto 2000 (Nexcelom Bioscience) with Acridine Orange (AO) / Propidium Iodide (PI) staining (viability: 99.7% for K562; 98.8% for 3T3). The cells were then combined at a 1:1 ratio to a final concentration of 2,000 cells/μl for PIP-seq capture. The mixed-species cell concentration was reverified using the Cellometer Auto 2000 (Nexcelom Bioscience) prior to single-cell capture.

#### DSP/Methanol and rehydration reagent preparation

1mg of DSP (ThermoFisher Scientific, Cat # A35393) was dissolved in 20ul of 100% Anhydrous DMSO (ThermoFisher Scientific, Cat # D12345). The 20uL of 50mg/ml DSP solution was then added to 800ul of ice cold 100% Methanol (Sigma-Aldrich, Cat # 34860) in a 1.5mL tube and gently mixed.

DSP/Methanol Rehydration buffer was made by adding to 893ul of nuclease-free water the following: 180ul of Saline sodium citrate (SSC) 20x (Sigma-Aldrich; CAT# S6639, 1.2ul of DTT 1M (Sigma-Aldrich; Cat # 646563), 6ul of Protector RNase Inhibitor 2,000U (Roche, 3335399001), 120ul of 10% BSA in DPBS (Sigma-Aldrich; Cat# A1595). DSP/Methanol/DEPC rehydration buffer was made similarly; however, instead of 6ul of Protector RNase inhibitor, 12ul of DEPC was used and 887ul of nuclease-free water.

#### Fresh vs DSP/Methanol PBMC sample preparation

Cryopreserved PBMC were thawed for 2min in 37°C water bath, washed twice using PBS containing 2% FBS, paletted using a swing bucket centrifuge (300 x g for 5 minutes, 4°C), then resuspended in 200ul Cell staining buffer (Biolegend, # 420201). Viability of samples were evaluated using Tryphan blue stain using Nexcelom Cellometer Auto 2000 (Nexcelom Bioscience) (97.7% viability). Samples were mixed well, split in two, then each incubated with TotalSeq-A hashtag antibodies (Biolegend) for 30minutes on ice. Cells were washed thrice with cell staining buffer then either resuspended in 820ul of freshly prepared chilled DSP/Methanol solution (added gradually with gentle swirling) or in Fluent cell resuspension buffer (FB0002440). DSP/Methanol sample was left on ice for 30minutes with occasional swirling then quenched with 20.4ul 1M Tris-HCL pH 7.5 (Invitrogen, Cat # 15567027). DSP/Methanol sample was then centrifuged (400 x g, 5min, 4°C) using bucket centrifuge, supernatant discarded, and resuspended in rehydration buffer. Both samples were filtered through 40um Flowmi cell strainers (Bel-Art, H13680-0040), counted with AO/PI stain using Nexcelom Cellometer Auto 2000 (Nexcelom Bioscience) and pooled to attain a 1:1 mixed ratio of both samples. Samples were further diluted using Fluent cell resuspension buffer (FB0002440), making up a concentration of 2000 cells/ul.

#### Multi-Fixation PBMC sample preparation

Cryopreserved PBMC were thawed for 2min in 37°C water bath, washed twice using PBS containing 2% FBS, paletted using a swing bucket centrifuge (300 x g for 5 minutes, 4°C), then resuspended in 600ul Cell staining buffer (Biolegend, # 420201). Viability of samples were evaluated using Tryphan blue stain using Nexcelom Cellometer Auto 2000 (Nexcelom Bioscience) (98% viability). Samples were mixed well, split into 6, with each incubated with TotalSeq-A hashtag antibodies (Biolegend) for 30minutes on ice. Cells were then washed thrice with cell staining buffer then either resuspended in 300ul of freshly prepared chilled DSP/Methanol solution (added gradually with gentle swirling),150ul Methanol (added gradually with gentle swirling), 150ul of fluent cell resuspension buffer (FB0002440) with methanol free to make 0.1% PFA solution (Life Technologies, Cat #28908),150ul of fluent resuspension buffer (FB0002440) PFA making 1% solution (Life Technologies, Cat #28908) or 150ul of fluent resuspension buffer (FB0002440). Methanol and the DSP/Methanol samples were left on ice for 30minutes with occasional swirling, and the DSP/Methanol sample was quenched with 40.8ul 1M Tris-HCL pH 7.5 (Invitrogen, Cat # 15567027) then split equally into two 150ul in new 1.5ml Eppendorfs. PFA samples were left at room temperature to fix for 5min then quenched with 1.25 M glycine solution (Life Technologies, J64365.30). All samples were centrifuged at 400 x g for 5 minutes at 4°C using bucket centrifuge, supernatant discarded, and resuspended in either fluent resuspension buffer (Fresh and PFA fixed samples) or rehydration buffer, with the DSP/Methanol/DEPC sample using a rehydration buffer that has DEPC (Sigma-Aldrich, Cat # D5758) instead of Protector RNase Inhibitor (Roche, 3335399001). All samples were filtered through 40um Flowmi cell strainers (Bel-Art, H13680-0040), counted with AO/PI stain (94% viability for fresh sample) using Nexcelom Cellometer Auto 2000 (Nexcelom Bioscience) and pooled with an attempt to attain equal ratio at a concentration of 3,000 cells/ul.

#### PIP-seq cell full length cDNA library prep

Cells were captured by adding 10 μl of sample (2,000 cells/ul K562:3T3 cells; 1650 cells/ul Fresh PBMCs, 2000 cells/ul DSP/Methanol vs Fresh PBMCs, 3000cells/ul Multi-Fix PBMC) directly into Fluent PIP bead reagent mix (PIPseq T20 Single Cell RNA kit v4.0). For K562:3T3 samples, and fresh only PBMC experiments, 2 μl of Recombinant RNase Inhibitor (Takara; cat no. 2313A) was also added. For fix vs fresh PBMC sample comparison, Protector RNase Inhibitor (Roche, 3335399001) was used. Cell capture, cell lysis, cDNA synthesis and whole transcriptome amplification (WTA) were performed as per the manufacturer’s instructions. Briefly, cells suspended in the emulsion were lysed and mRNA was captured on PIP-beads before breaking the emulsion prior to reverse transcription. cDNA underwent WTA, using 11 cycles for 3T3:K562 cDNA and 14 cycles for PBMC cDNA’s, with 8 min extensions. Amplified cDNA was isolated from the PIPs then SPRI purified and quantified using Qubit. Library integrity assessment was performed on a high-sensitivity D5000 ScreenTape on 4150 Tapestation (Agilent Technologies).

#### PIP-seq Illumina Library prep and sequencing

Libraries were prepared for Illumina sequencing as per the manufacturer’s instructions (PIPseq T20 Single Cell RNA kit v4.0). Briefly, 30ng of PBMC cDNA and 100 ng of K562:3T3 cDNA were input into the PIP-seq library prep module to undergo fragmentation, end repair, A-tailing, and adapter ligation and underwent indexing PCR using 11 cycles for PMBC and 9 cycles for K562:3T3. Final libraries were quantified using Qubit and fragment sizes were verified using a high-sensitivity D1000 ScreenTape on 4150 Tapestation (Agilent Technologies) prior to equimolar pooling and sequencing at BCL on an either an Illumina NovaSeq X (28-bp Read 1, 10-bp i7 index, 10-bp i5 index, and 90-bp Read 2), NovaSeq SP (54-bp Read 1, 8-bp i7 index, 8-bp i5 index, and 67-bp Read 2) or NovaSeq 2000 (28-bp Read 1, 10-bp i7 index, 10-bp i5 index, and 90-bp Read 2).

#### PIP-seq cell Hashtag library prep and sequencing

Following WTA cDNA of PIP-seq captures, the supernatant of first SPRI cleanup which contained Hashtag cDNA was further processed as per manufacturer’s instructions. Briefly, following another round of SPRI purification, libraries were amplified (8 cycles) using primers that appended illumina adapters. Libraries were SPRI cleaned, evaluated using high-sensitivity D1000 ScreenTape on 4150 Tapestation (Agilent Technologies) and sequenced either on Illumina NovaSeq SP (54-bp Read 1, 8-bp i7 index, 8-bp i5 index, and 67-bp Read 2) or NovaSeq 2000 (28-bp Read 1, 10-bp i7 index, 10-bp i5 index, and 90-bp Read 2).

#### 10x Genomics full length cDNA library prep

Using single cell 3′ V3.1 (10x Genomics, 1000269) and single Cell 5′ v2 (10x Genomics, 1000267) kits, captures were carried out according to the manufacturer’s protocol by Broad Clinical Labs on two 10x chromium controllers simultaneously (10x Genomics, 120270). Briefly, cell suspensions loaded onto the Chromium Next GEM Chip G along with gel beads and partitioning oil. Following GEM generation, samples were immediately subjected to reverse transcription using a thermal cycler with the following conditions: 53°C for 45 minutes, 85°C for 5 minutes, and a hold at 4°C. After RT, emulsions were broken, and cellular debris was removed via a Silane bead cleanup (10x Genomics, 2000048). The purified cDNA was amplified under the following PCR conditions: initial denaturation at 98°C for 3 minutes, followed by 12 cycles of 98°C for 15 seconds (denaturation), 63°C for 20 seconds (annealing), and 72°C for 1 minute (extension). The amplified cDNA underwent a 0.6× SPRI bead cleanup (Beckman Coulter) and was quantified using a Qubit fluorometer (Thermo Fisher Scientific). The final full-length cDNA was then divided into aliquots for short-read and long-read library constructions.

#### 10x Genomics Illumina library construction

Library construction was performed using an automated workflow. The cDNA was fragmented and repaired with the addition of an A-tail by incubating at 32°C for 5 minutes and 65°C for 30 minutes, using the Chromium Single Cell 3′ Library Construction Kit v3.1 (10x Genomics, PN-2000160). A 0.8× double-sided SPRI bead cleanup (Beckman Coulter) was performed to remove undesired fragments. Adapters were ligated onto the DNA at 20°C for 15 minutes, followed by another 0.8× SPRI bead cleanup. Dual-indexed barcodes (10x Genomics, 1000215) were added, and libraries were amplified using PCR with the following conditions: initial denaturation at 98°C for 45 seconds, followed by 12 cycles of 98°C for 20 seconds (denaturation), 54°C for 30 seconds (annealing), and 72°C for 20 seconds (extension). Libraries underwent a final 0.8× double-sided SPRI bead cleanup for size selection. The quality of the final libraries was assessed using Quant-iT PicoGreen (Thermo Fisher Scientific) and an Agilent TapeStation (Agilent) to confirm the expected library size. Libraries were pooled to achieve the desired number of reads per cell for downstream analysis. Libraries were sequenced using NovaSeqX (28-bp Read 1, 10-bp i7 index, 10-bp i5 index, and 90-bp Read 2).

### Long-read sample preparation

#### ISO-seq (1-mer) library prep for PacBio sequencing

3.2 ng of cDNA was amplified in a 100 μl *MAS-*PCR reaction using the following reaction conditions: 50 μl of MAS PCR mix, 10 μl of custom indexed Ai/Qi MAS primers (see Supplementary Table 8 for MAS primers; to enable multiplexing), 15 ng of cDNA library, adding H2O to bring final volume to 100 ul. Reactions were amplified using the following cycling conditions: 98°C for 3 minutes, followed by 8 cycles of 98°C for 20 s, 68°C for 30 s and 72°C for 8 minutes, followed by a final 72°C extension for 10 minutes. The amplified libraries were cleaned using a 0.8× SPRIselect beads (Beckman Coulter B23318), eluted into 60 μl elution buffer, and quantified with Qubit (Thermo Fisher Scientific, Q32851). Barbell formation was carried out by mixing 500 ng of MAS PCR product in 55 μl volume with 2 μl MAS enzyme and incubated at 37°C for 30 minutes. Subsequently, 3ul of barcode MAS adapter and 20 μl of MAS ligation additive was mixed into the sample followed by 10 μl of MAS ligase buffer and 10 μl fo MAS ligase. Sample was thoroughly mixed by gentle pipetting using a wide bore pipette tip to avoid bubble formation. MAS array formation reactions were incubated at 42°C for 60 minutes followed by 4°C hold (52°C lid). Array formation reactions were cleaned using 0.8× SPRIselect beads (Beckman Coulter B23318), eluted into 43 μl elution buffer and quantified with Qubit (Thermo Fisher Scientific, Q32851). DNA Damage Repair was carried out by mixing 42 μl of sample with 6 μl of Repair buffer and 2 μl of DNA repair mix, and incubated at 37°C for 30 minutes (lid 47°C). DNA damage repair reactions were cleaned using 0.8× SPRIselect beads (Beckman Coulter B23318), eluted into 40 μl elution buffer and quantified with Qubit (Thermo Fisher Scientific, Q32851).

Nuclease treatment was carried out by mixing 40 μl of DNA damage repaired library with 5 μl Nucelase buffer and 5 μl Nuclease mix and incubated at 37°C for 60 minutes followed by 4°C hold (lid 47°C). The final MAS array libraries were cleaned using 0.8× SPRIselect beads (Beckman Coulter B23318), eluted into 20 μl elution buffer and quantified with Qubit (Thermo Fisher Scientific, Q32851). Library size was obtained using a Genomic DNA ScreenTape and 4150 Tapestation (Agilent Technologies). Multiplexed ISO-seq libraries were pooled in equimolar amounts and sequenced on two Revio SMRT cells (PN: 102-202-200).

#### TSO artifact purification

TSO artifact removal was carried out as per manufacturer’s instructions. Briefly, 15 ng of cDNA from 3′ 10x PBMC and 5′ 10x PBMC were each used for TSO PCR using the following reaction conditions: 25 μl of MAS PCR mix, 10 μl of MAS 5′ or 3′ capture primer mix, 15 ng of 10x 3′ or 10× 5′ cDNA library, adding H2O to bring final volume to 50 ul. Reactions were amplified using the following cycling conditions: 98°C for 3 minutes, followed by 5 cycles of 98°C for 20 s, 68°C for 30 s and 72°C for 8 minutes, followed by a final 72°C extension for 10 minutes. The amplified libraries were cleaned using 0.8× SPRIselect beads (Beckman Coulter B23318) and quantified with Qubit (Thermo Fisher Scientific, Q32851). They were then further purified with 10 μl MAS capture beads. Following the binding and washing steps, the beads were reconstituted in 40 μl of elution buffer. 2 μl of MAS enzyme was added to each sample and incubated at 37°C for 30 minutes to cleave the captured DNA from the MAS capture beads. Eluant was moved to a fresh PCR tube and cleaned using a 0.8× SPRIselect beads (Beckman Coulter B23318), eluted into 45 μl of elution buffer and quantified with Qubit (Thermo Fisher Scientific, Q32851). This product was then used as input into MAS PCR.

#### MAS-seq 16-mer indexed library prep for PacBio sequencing

The following samples were prepared for indexed Kinnex 16mer workflow:

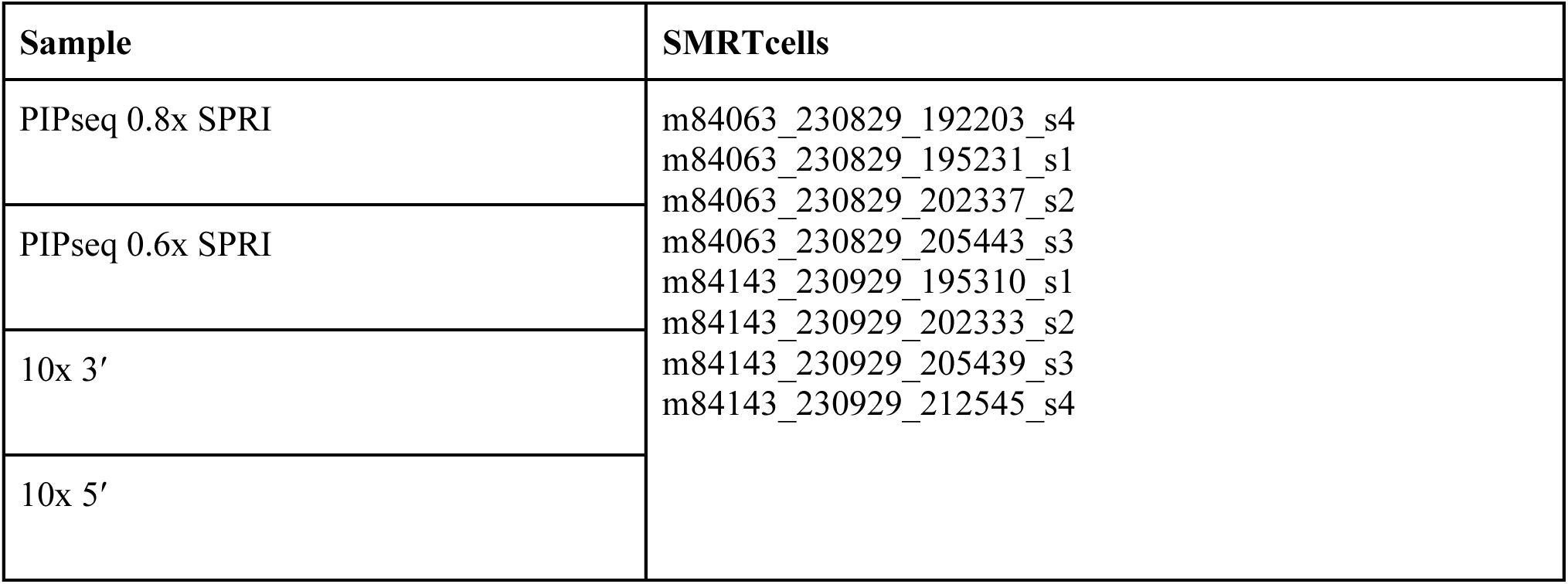

MAS-seq 16-mer libraries were prepared according to manufacturer’s instructions (PacBio MAS-seq kit Cat, 102-407-900). Briefly, 55ng cDNA was split between 16 MAS PCR reactions, each with the following final reaction conditions: 50 μl of MAS PCR mix, 15 ng of cDNA library, adding H2O to bring final volume to 90 ul, then adding 10 μl of of MAS primer mix pairs composed of sequential primers to each respective PCR reaction (see primer table). To generate indexed arrays, we substituted indexed primers for the A and Q primers (see primer table). Reactions were amplified using the following cycling conditions: 98°C for 3 minutes, followed by 8 cycles of 98°C for 20 s, 68°C for 30 s and 72°C for 8 minutes, followed by a final 72°C extension for 10 minutes. The amplified libraries were pooled and cleaned using 0.8× SPRIselect beads (Beckman Coulter B23318), eluted into 50 μl elution buffer and quantified with Qubit (Thermo Fisher Scientific, Q32851). MAS array formation was carried out by mixing 4 ug of MAS PCR product in 47 μl volume with 10 μl MAS enzyme and incubated at 37°C for 30 minutes. Subsequently, 3ul of barcode MAS adapter and 20 μl of MAS ligation additive was mixed into the sample followed by 10 μl of MAS ligase buffer and 10 μl of MAS ligase. Sample was thoroughly mixed by gentle pipetting using a wide bore pipette tip to avoid bubble formation. MAS array formation reactions were incubated at 42°C for 60 minutes followed by 4°C hold (52°C lid). Array formation reactions were cleaned using 0.8× SPRIselect beads (Beckman Coulter B23318), eluted into 43 μl elution buffer and quantified with Qubit (Thermo Fisher Scientific, Q32851). DNA Damage Repair was carried out by mixing 42 μl of sample with 6 μl of Repair buffer and 2 μl of DNA repair mix, and incubated at 37°C for 30 minutes (lid 47°C). DNA damage repair reactions were cleaned using 0.8× SPRIselect beads (Beckman Coulter B23318), eluted into 40 μl elution buffer and quantified with Qubit (Thermo Fisher Scientific, Q32851). Nuclease treatment was carried out by mixing 40 μl of DNA damage repaired library with 5 μl Nucelase buffer and 5 μl Nuclease mix and incubated at 37°C for 60 minutes followed by 4°C hold (lid 47°C). The final MAS array libraries were cleaned using 0.8× SPRIselect beads (Beckman Coulter B23318), eluted into 20 μl elution buffer and quantified with Qubit (Thermo Fisher Scientific, Q32851). Library size was obtained using a Genomic DNA ScreenTape and 4150 Tapestation (Agilent Technologies).

#### MAS-seq 16-mer library prep for PacBio sequencing

The following samples were prepared for standard Kinnex 16mer workflow:

| Sample | SMRTcells |
| --- | --- |
| Fixation protocol PBMC experiment | m84175_240224_082414_s4.hifi_reads.bcM0004.bam |
| Second round fix/fresh comparison | m84250_240530_044215_s1.hifi_reads.bcM0001.bam<br>m84250_240530_064125_s2.hifi_reads.bcM0001.bam |
| 10x 3', no cleanup | m84252_240709_213425_s4.hifi_reads.bcM0002.bam |
| 10x 3', with cleanup | m84252_240709_213425_s4.hifi_reads.bcM0003.bam |
| 10x 3dG | m84252_240709_213425_s4.hifi_reads.bcM0004.bam |

MAS-seq 16-mer libraries were prepared according to manufacturer’s instructions (PacBio MAS-seq kit Cat, 102-407-900). Briefly, 55ng cDNA was split between 16 MAS PCR reactions, each with the following final reaction conditions: 50 μl of MAS PCR mix, 15 ng of cDNA library, adding H2O to bring final volume to 90 ul, then adding 10 μl of of MAS primer mix pairs composed of sequential primers to each respective PCR reaction (see primer table). Reactions were amplified using the following cycling conditions: 98°C for 3 minutes, followed by 8 cycles of 98°C for 20 s, 68°C for 30 s and 72°C for 8 minutes, followed by a final 72°C extension for 10 minutes. The amplified libraries were pooled and cleaned using 0.8× SPRIselect beads (Beckman Coulter B23318), eluted into 50 μl elution buffer and quantified with Qubit (Thermo Fisher Scientific, Q32851). MAS array formation was carried out by mixing 4 ug of MAS PCR product in 47 μl volume with 10 μl MAS enzyme and incubated at 37°C for 30 minutes. Subsequently, 3ul of barcode MAS adapter and 20 μl of MAS ligation additive was mixed into the sample followed by 10 μl of MAS ligase buffer and 10 μl of MAS ligase. Sample was thoroughly mixed by gentle pipetting using a wide bore pipette tip to avoid bubble formation. MAS array formation reactions were incubated at 42°C for 60 minutes followed by 4°C hold (52°C lid). Array formation reactions were cleaned using 0.8× SPRIselect beads (Beckman Coulter B23318), eluted into 43 μl elution buffer and quantified with Qubit (Thermo Fisher Scientific, Q32851). DNA Damage Repair was carried out by mixing 42 μl of sample with 6 μl of Repair buffer and 2 μl of DNA repair mix, and incubated at 37°C for 30 minutes (lid 47°C). DNA damage repair reactions were cleaned using 0.8× SPRIselect beads (Beckman Coulter B23318), eluted into 40 μl elution buffer and quantified with Qubit (Thermo Fisher Scientific, Q32851). Nuclease treatment was carried out by mixing 40 μl of DNA damage repaired library with 5 μl Nucelase buffer and 5 μl Nuclease mix and incubated at 37°C for 60 minutes followed by 4°C hold (lid 47°C). The final MAS array libraries were cleaned using 0.8× SPRIselect beads (Beckman Coulter B23318), eluted into 20 μl elution buffer and quantified with Qubit (Thermo Fisher Scientific, Q32851). Library size was obtained using a Genomic DNA ScreenTape and 4150 Tapestation (Agilent Technologies).

#### 3′-Deoxyguanosine (3dG) Template Switch Oligo Experiment

The 3′-Deoxyguanosine (3dG) template switch oligo was obtained from Biosyn Corporation, sequence: /5BiosG/AAGCAGTGGTATCAACGCAGAGTACATrGrG{3dG} with RNAse free HPLC purification. Following the standard 10x protocol, the 3dG TSO was resuspended to 1mM in elution buffer. The TSO comparison study was performed using PBMCs (AllCells) on the 10x 3′ Gene Expression Kit (10x Genomics, Cat #PN-1000269). The PBMC samples were run in parallel comparing following the standard protocol and a 3dG protocol with the only modification being the addition of 2.4ul of 1mM 3dG TSO in place of the standard TSO.

### Computational Methods

#### Code Availability

The code to perform the computational analysis for this paper is available at www.github.com/MethodsDev/mdl-sc-isoform-paper. This repository contains Jupyter notebooks that document all of the analysis performed in this manuscript and produce all of the figures. In addition, the analysis relies on the following custom packages available on Github:

| Repository | URL | Description |
| --- | --- | --- |
| `marti` | https://github.com/PopicLab/marti | Artifact analysis framework (C++) |
| `callao` | [MethodsDev/callao](https://github.com/MethodsDev/callao) | Tool for demultiplexing $A_i$ / $Q_i$ indexed libraries |
| `bouncer` | [MethodsDev/bouncer](https://github.com/MethodsDev/bouncer) | Python package for matching scRNAseq barcodes |
| `barcode-symspell` | [MethodsDev/barcode-symspell](https://github.com/MethodsDev/barcode-symspell) | Efficient edit-distance library used by `bouncer` |
| `isoscelles` | [MethodsDev/isoscelles](https://github.com/MethodsDev/isoscelles) | Python package for single-cell isoform analysis |
| `bugzapper` | [MethodsDev/bugzapper](https://github.com/MethodsDev/bugzapper) | Tool for constructing a barcode-UMI-gene array from a BAM file, for subsampling |
| `mdl-core` | [MethodsDev/mdl-core](https://github.com/MethodsDev/mdl-core) | Python package of basic utilities used by other tools. |
| `ITV` | [MethodsDev/ITV](https://github.com/MethodsDev/ITV) | Integrative Transcriptomics Viewer, for visualization |

#### References

The following human references were used:

– FASTA: GRCh38_no_alt genome
– BED (provided to minimap2): Gencode v39 annotation
– GTF: Gencode v39 basic annotation – only transcripts with Gencode’s basic tag
– Gencode polyA site annotation
– polyA motif file (distributed by SQANTI3, from Beaudoing *et al.*, 2000)

For the barnyard experiment, we used the mouse-and-human genome reference available from 10x and built a new STAR index that was compatible with PIPseeker.

#### Short-read sequencing data processing

Data from the Illumina NovaSeq X was demultiplexed with bcl2fastq2 (v2.20.0.422). 10x 3′ and 10x 5′ experiments were then processed using cellranger (v7.1.0) using the reference genome and GTF specified above. The PIPseq samples were processed with PIPseeker (PBMC sample: v2.1.4; Initial fixation experiments: v3.0.5; Second round of fixation experiments: v3.3.0). The PBMC sample was aligned to the human reference and the barnyard experiment was aligned to a combined human and mouse reference, derived from the 2020-A reference made available by 10x Genomics at https://www.10xgenomics.com/support/software/cell-ranger/downloads#reference-downloads.

#### Long-read sequencing data processing

Long-read sequencing was performed on two instruments: monomer cDNA sequencing libraries were sequenced on the PacBio Sequel IIe, while MASseq arrays were sequenced on PacBio Revio sequencers.

Sequel IIe samples were demultiplexed by first annotating primers with lima (v2.9.0) and then separating individual samples with a custom tool, named callao. callao uses the annotations created by lima and outputs one BAM for each sample. For the analysis of artifacts in these data, we included reads that contained the same index on either end (A_i_-A_i_ and Q_i_-Q_i_ reads), which are produced when cDNA has the same adapter on either end. In MASseq arrays these artifacts can produce palindromic arrays of different lengths (e.g. ABCBA).

Revio samples were demultiplexed in the same fashion and then deconcatenated using skera (v1.2.0), using custom array lists based on the A_i_ and Q_i_ adapters that correspond to the indexes used.

After the demultiplexing and deconcatentation of MASseq arrays, reads were classified using marti, a tool we developed for in-depth artifact analysis. At a high level, marti classifies each read based on the presence, absence, and location of predefined target sequence features (such as, PCR and sequencing adapters) within the read. It takes as input an unaligned BAM file and a YAML configuration file with search parameters (e.g., the expected sequencing error rate and minimum poly(A) tail length) and target sequences of interest. It outputs (1) a BAM file with annotated cDNA reads, where per-read annotations include the artifact category, the coordinates of all the target sequence features and the native cDNA sequence, and the structural representation of the read (which captures the order and orientation of the target features within the read), and (2) a report of counts for each artifact category and read structure. marti classifies the reads into the following classes: (1) *Proper*: non-artifactual read, (2) *TSO-TSO*: RT artifact caused by the internal priming of the switch-leader-sequence (SLS, terminal 3′ end of the TSO which is not part of the TSO adapter sequence) of the TSO, characterized by the presence of a TSO at both ends of the reads, (3) *RT-RT*: RT artifact caused by the internal priming of the oligo dT (from the RT primer), characterized by the presence of the RT PCR adapter and a poly(A) at both ends of the read, (4) *internal-priming-RT-adapter*: PCR artifact caused by the internal priming of the RT PCR adapter, (5) *internal-priming-TSO-adapter*: PCR artifact caused by the internal priming of the TSO PCR adapter, (6) *only-polyA*: artifact class assigned when only As or Ts are found inside the read sequence, likely a result of RNA degradation, (7) *missing-adapter*: sequencing artifact, assigned when the read is missing at least one PCR adapter at either end, (8) *internal-adapter*: sequencing artifact, assigned when an extra adapter is found inside the read, (9) *retained-SMRTbell*: sequencing artifact in PacBio reads, assigned when the PacBio SMRTbell adapter is found inside the read, (10) *unknown*: artifact of unknown type assigned when the read structure does not correspond to a proper read nor any other known artifact classes, which allows users to identify novel artifact classes (by examining the discovered read structures) and filter out such constructs from downstream analysis. Since some artifact classes may correspond to multiple read structures, marti separately reports and quantifies each observed read structure within each artifact class. For instance, the *internal-priming-TSO-adapter* artifact can be detected in the following three different read configurations (in each orientation): (TSO PCR adapter, cDNA, poly(A), reverse_complement(RT PCR adapter), (TSO PCR adapter, cDNA, reverse_complement(TSO)), and (TSO PCR adapter, cDNA, reverse_complement(TSO PCR adapter). With the exception of *internal polyA priming*, Supplementary Figure 6 summarizes the artifact classes and read structures reported by marti.

#### Short-read analysis

##### Barnyard data

Publicly available data for human-mouse mixture experiments were obtained from https://www.10xgenomics.com/datasets. We downloaded the unfiltered raw h5 matrix files for two barnyard experiments, which had been aligned to the same human-mouse combined reference used for the PIPseq experiment:

– 10k_hgmm_3p_nextgem_Chromium_Controller_raw_feature_bc_matrix.h5 for 10x 3′ v3.1 data
– 10k_hgmm_5pv2_nextgem_Chromium_Controller_raw_feature_bc_matrix.h5 for 10x 5′ v2 data

We called cells and doublets using a UMI threshold adjusted for the relative depth of the three experiments. To calculate doublet rates relative to throughput, 10x 3′ cells were called at 2,750 UMIs/barcode, 10x 5′ cells were called at 3,000 UMIs/barcode and PIPseq cells were called at 10,000 UMIs/barcode. To measure ambient levels in these experiments we considered cells with ≥10,000 UMIs of one species and <1,000 UMIs of the other as confident single-species barcodes, and considered the percentage of cross-species UMIs as a proxy for ambient contamination.

##### PBMC data

As a preliminary analysis, each dataset was analyzed separately using all reads. For the 10x data we filtered to barcodes with >1,000 UMIs. After estimating the number of cells in each sample, we subsampled the full sequencing data so that each dataset contained roughly 100,000 reads per cell and selected barcodes with >1,000 UMIs after subsampling (Supplementary Fig. 13). All downstream analysis was performed on the subsampled data.

Gene selection and clustering were performed as described in J. Langlieb & N.S. Sachdev, 2023. Briefly, we used a binomial model to estimate the expected rate of non-zero counts in the case of homogenous gene expression in a population. A gene was selected as variable if the observed percentage of cells with non-zero count deviated from Poisson expectation by more than a specified percentage. A minimum of 5% deviation from expectation was used for gene selection during clustering.

Clustering was performed on each dataset separately, with a hierarchical strategy^1^. Each round of clustering consisted of four steps. First, gene selection was performed based on the above binomial model. Next, counts were square-root transformed. Third, *k*-nearest neighbor (kNN using cosine distance, with *k* = 80) and shared nearest neighbor (SNN) graphs were constructed. Finally, the SNN graph was clustered using the Leiden algorithm over a range of resolution parameters to find the lowest resolution that yielded multiple clusters^2^. The resulting clusters were each recursively re-clustered with the same strategy, and the process was continued until no resolution parameter yielded multiple clusters, or the resulting clusters were too small to compute a meaningful SNN graph (if *n* < *k* = 80 cells, the graph is fully connected and all edge weights are equal). Clusters were labeled by manual inspection, using a set of canonical markers for peripheral blood cells.

For visualization purposes we used a force-directed graph embedding based on the same type of shared nearest neighbors graph. In this case a minimum deviation of 10% was used. The SFDP layout algorithm was run on the SNN graph and the three embeddings were rotated and reflected as needed to align similar cell populations to each other.

##### Hashtagged Fixation Comparisons

Count matrices of cells and hashtags were generated using PIPseeker’s built-in functionality. We selected the highest-sensitivity filtered matrix (level 5, in PIPseeker’s output) for processing and took the barcodes called for each hashtag for further analysis. Counts were processed using Seurat v4.3 using standard single-cell analysis practices and parameters (dim 1:15, res = 0.8). Hashtag counts were CLR normalised and cells were demultiplexed to their respective fixation or fresh using Seurat inbuilt *HTODemux* function (0.99 positive quantile).

### Annotation of Cell types

Canonical markers commonly used for differentiating cell types were used for annotation of clusters (PMCID: PMC8238499). First cells were demarcated by broader lineage associated markers (Myeloid:*LYZ*, B-cell:*MS4A1*, T-cell/NK/ILC:*CD3*, Red blood cell:*PPBP*), then clusters within each lineage was further stratified as per following: Classical Monocyte: *CD14+FCGR3A-,* Non-classical monocyte: *CD14*+ *FCGR3A+*, DC: *CLEC10A+* / *CD1C+ / IRF7+*, CD4 T-cells: *CD4*+*CD8*-, CD8 T-cells: *CD8*+*CD4*-*NCAM1*-, NK: *TRBC1-XCL1*+*NCAM1*+, CTL: CD8+/− GNLY+,*TRBC1*+/− ILC: TRDC+ *CD8*+/− *NCAM1*+/− *TRBC1+/−,* Naïve T-cell: *CD3*+ *SELL+ CCR7+*

### Long-read analysis

#### Barcode extraction

To extract barcodes and UMIs from long-read data we extracted valid barcodes and UMIs from the marti-annotated sequences. Only Proper reads were used for alignment and isoform identification. Given the expected single-cell library structure and location of the adapters (provided by marti annotation), we extracted the putative location of the cell barcode and matched the region against a whitelist of known barcodes (Supplementary Fig. 14). For matching we used a maximum edit distance of 1 and only accepted barcodes with a single unambiguous match. After the barcode has been located, the adjacent sequence of the appropriate length is taken as the UMI. Using the structure of the read we also extracted the cDNA and reoriented all reads to the forward orientation for mapping.

To perform barcode matching on very long (39bp) PIPseq barcodes we developed a new library, bouncer, which uses a symmetric-deletion index for efficient lookup of match candidates. This is an extension of the SymSpell method first developed for spell-checking by Wolfe Garbe (https://github.com/wolfgarbe/SymSpell). The most significant change from the original algorithm is the use of a partitioning method to efficiently store the very large (85M barcode) whitelist of 39bp PIPseq barcodes. The code for bouncer and the SymSpell implementation barcode-symspell are available on GitHub (see above).

#### Read alignment and filtering

After extracting cDNA and orienting all reads to the forward strand, we aligned reads to the GRCh38 reference genome using minimap2 (v2.28)^3^. Alignment was performed with the following settings:

minimap2 –t 16 –ayx splice:hq –uf –G1250k –Y –-MD –-junc-bed [gencode_v39_bed] [GRCh38_fasta] [input_fastq] > [output]

Of note is –G1250k to allow introns up to the maximum size annotated in the human genome, and –uf to only perform alignment on the forward strand of the input, as we know the original orientation of the cDNA from our annotation. While we use the “basic” Gencode annotation for isoform quantification, here we use the full annotation to give minimap2 access to additional putative splice sites. After alignment, we filtered the aligned BAMs for high-scoring primary alignments. High-scoring is defined as: MAPQ ≥ 60 or [alignment score] / [query len] > 0.9. In our testing, including secondary and supplemental alignments did not improve the isoform identification for this dataset, and their presence complicated the internal priming analysis.

#### Isoform identification

IsoQuant (v3.4.2) was run on the filtered primary reads with the GRCh38 genome, the Gencode v39 basic reference, and the following options: –-complete_genedb, –-stranded forward to indicate the reads are already correctly oriented, and –-polya_requirement never because we have removed the poly(A) tails from the reads^4^. We found that including poly(A) tails during alignment led to more reads misaligned to pseudo-genes in the genome.

To compare the results of IsoQuant across the experiments, we used gffcompare (v0.12.6) to match the novel transcripts in the transcript_models.gtf output by IsoQuant to the input reference, using the –-strict-match option^5^. This comparison was restricted to transcripts with at least five assigned reads.

#### Priming analysis and read annotation

We categorized each read according to the genomic context of their alignment, and assigned them to one of 19 categories, of which four categories were considered to be expected priming events. To be classified as an expected priming event, a read must terminate near a known transcription stop site or be adjacent to a poly(A) signal motif. If the read does not terminate near an annotated stop site, it must not contain a “genomic poly(A)”, defined as no more than 12 adenines and no more than 6 in a row starting within the 20 bases downstream of the read. These thresholds were determined empirically from analysis of bulk and single-cell mRNA datasets and are similar to the thresholds used by Pacific Biosciences’ pigeon tool^6^.

The output from the priming analysis is a set of annotated BAM files, with metadata for each read indicating its priming analysis as well as additional metadata from IsoQuant, including the transcript assignment and the SQANTI classification (e.g. full or incomplete splice match) for each read. Having produced these annotated BAMs we could efficiently quantify the number of transcripts in different categories (known transcripts, novel-in-catalog transcripts from known genes, and novel not-in-catalog transcripts from known and novel genes) and calculate the proportion of reads for each transcript that came from expected or unexpected priming sites.

#### Coverage analysis

After creation of the annotated BAM files, coverage could be efficiently calculated for each transcript by filtering the tagged reads based on the annotated tags. To construct an average profile, the coverage for each transcript was scaled to an equal number of bins (n = 1000) and the coverage was interpolated over that range. This allowed us to create an average profile over transcripts of differing lengths, either within a certain range of lengths or over the entire observed transcriptome.

## Acknowledgements

This work was supported both by Broad Clinical Labs and a collaboration agreement between Pacific Biosciences of California, Inc. and the Broad Institute.

## Supplementary Figures

**Supplementary Figure 1:**
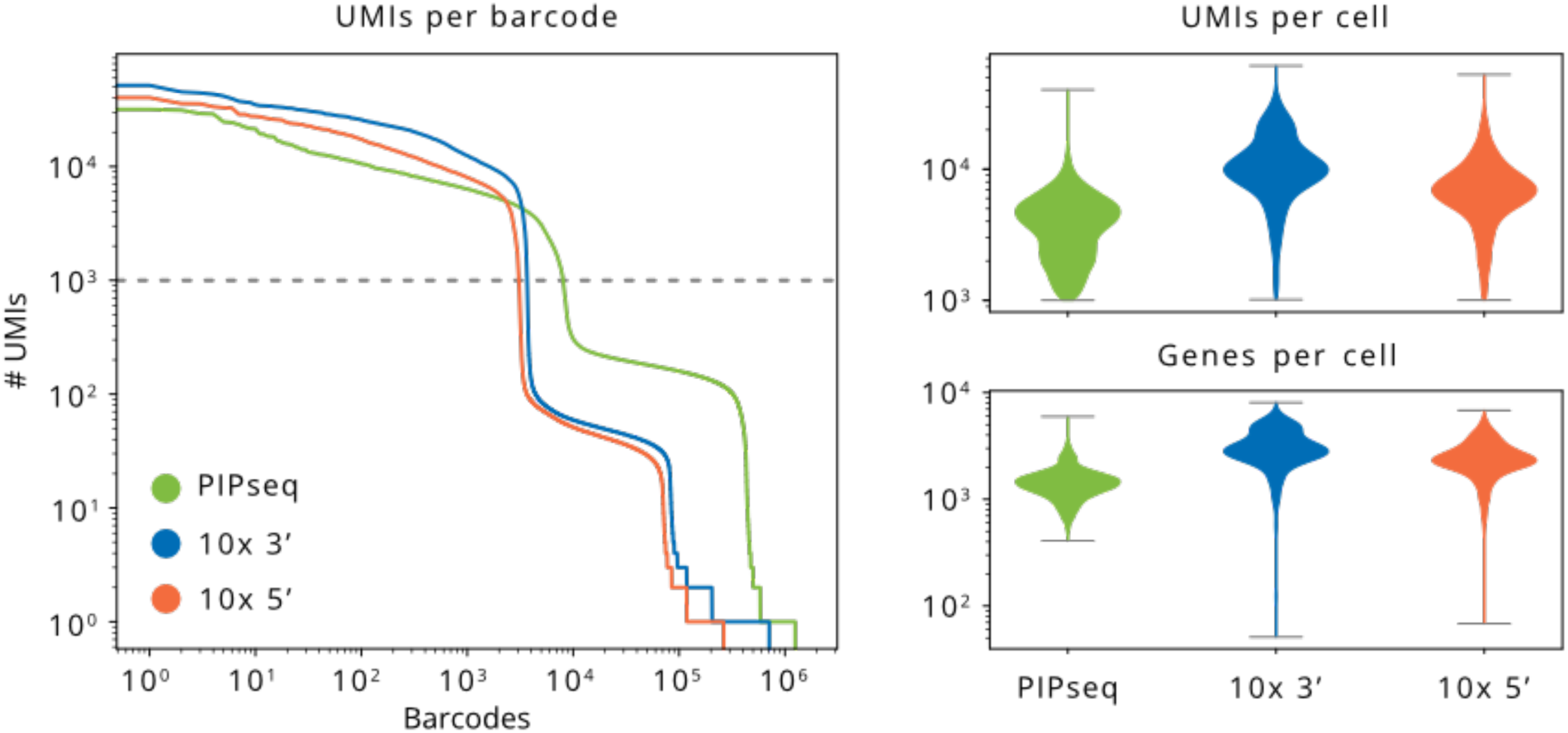
Kneeplots and metrics for the short-read libraries, subsampled to ∼100,000 reads per cell.

**Supplementary Figure 2:**
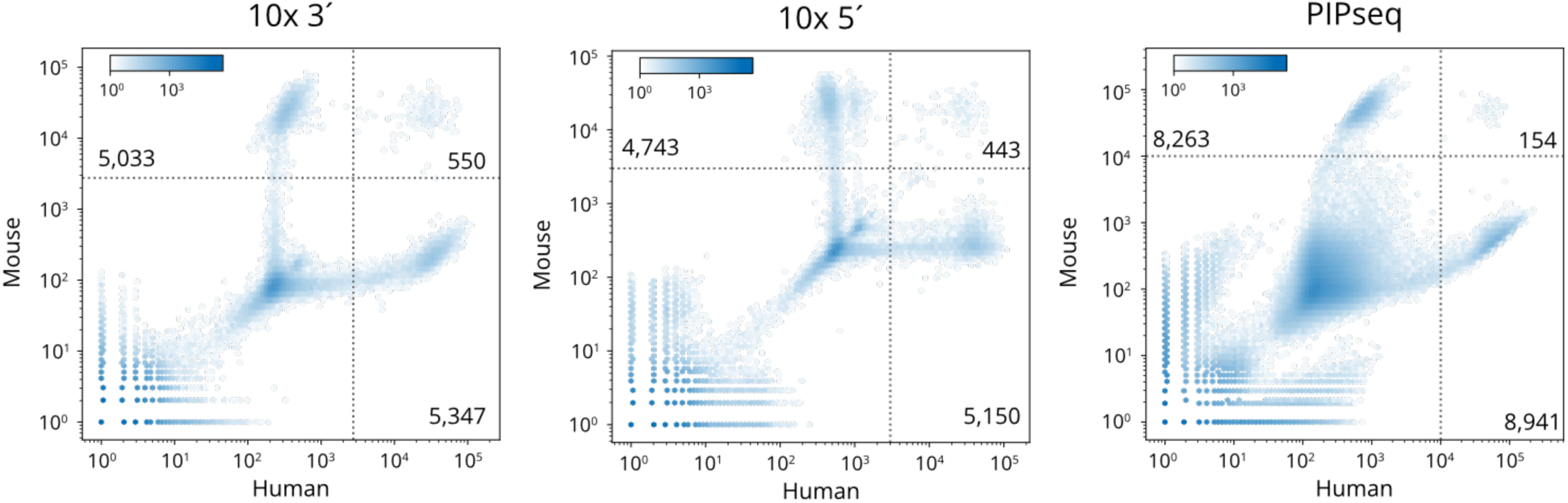
Barnyard data showing the number of cells and doublets called for each experiment, as well as the distribution of cross-species contamination from ambient mRNA. 10x 3′ and 5′ data is from public datasets made available by 10x Genomics. The top left quadrant contains mouse cells, while the bottom right quadrant contains human cells. The top right quadrant contains doublets of mouse and human (10x 3′:; 10x 5′:; PIPseq:). Note that the expected doublet count is twice this number, as we cannot detect doublets of the same species.

**Supplementary Figure 3:**
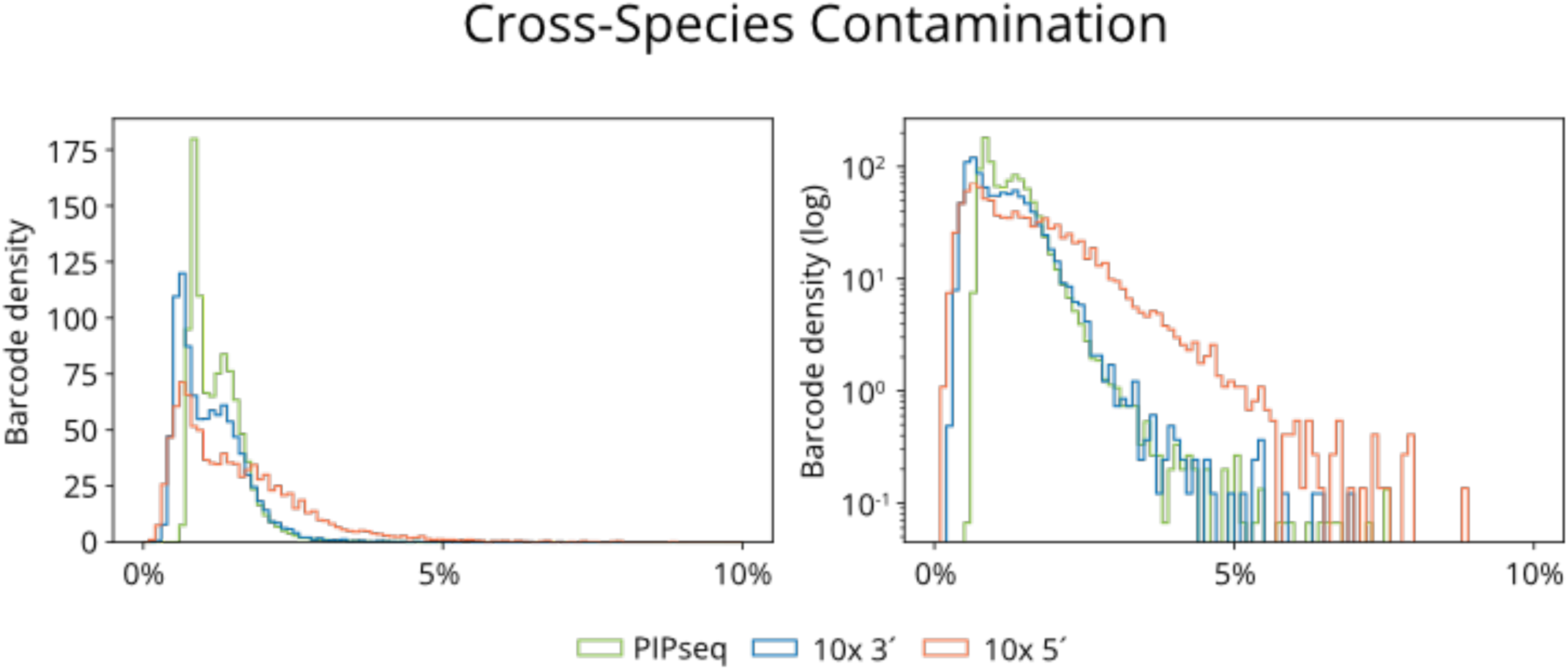
Distribution of ambient contamination (measured via cross-species alignment) estimated from barnyard experiment data, based on barcodes called confidently as one species (>10,000 UMIs) but not the other (<1,000 UMIs from the other species). On the right the distribution is log-scaled to show outliers.

**Supplementary Figure 4:**
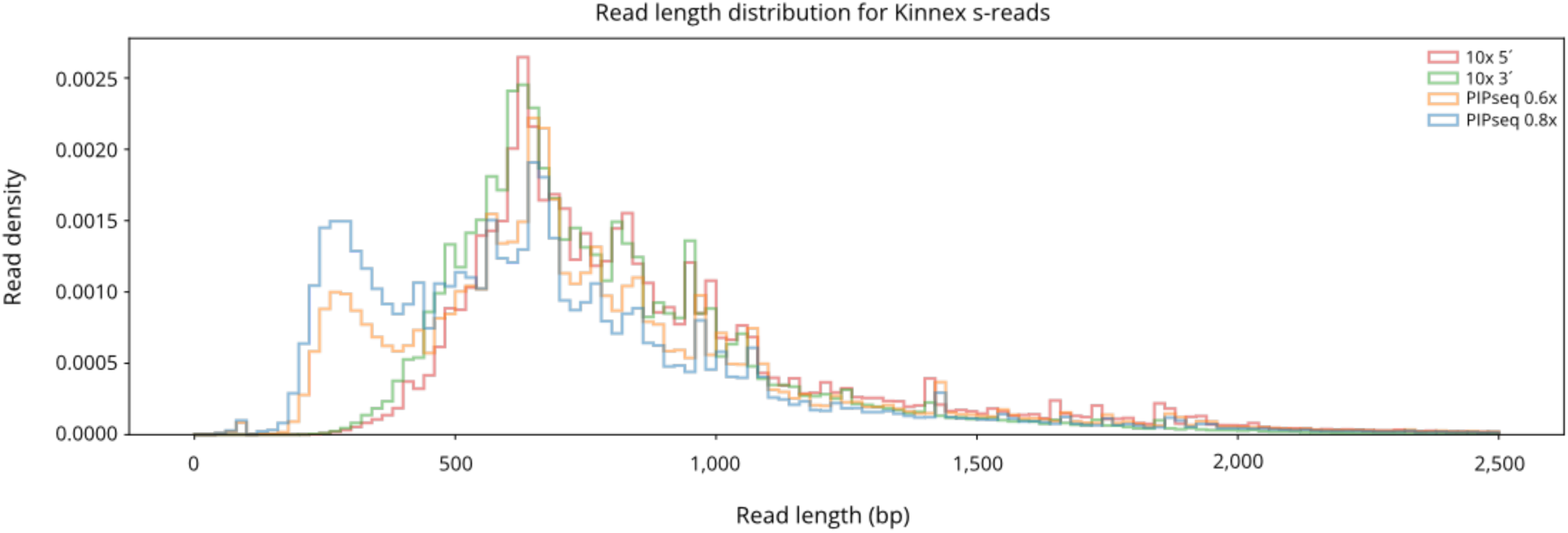
Read length distributions for Kinnex experiments. 0.8x and 0.6x refer to SPRI ratios used for PIPseq.

**Supplementary Figure 5:**
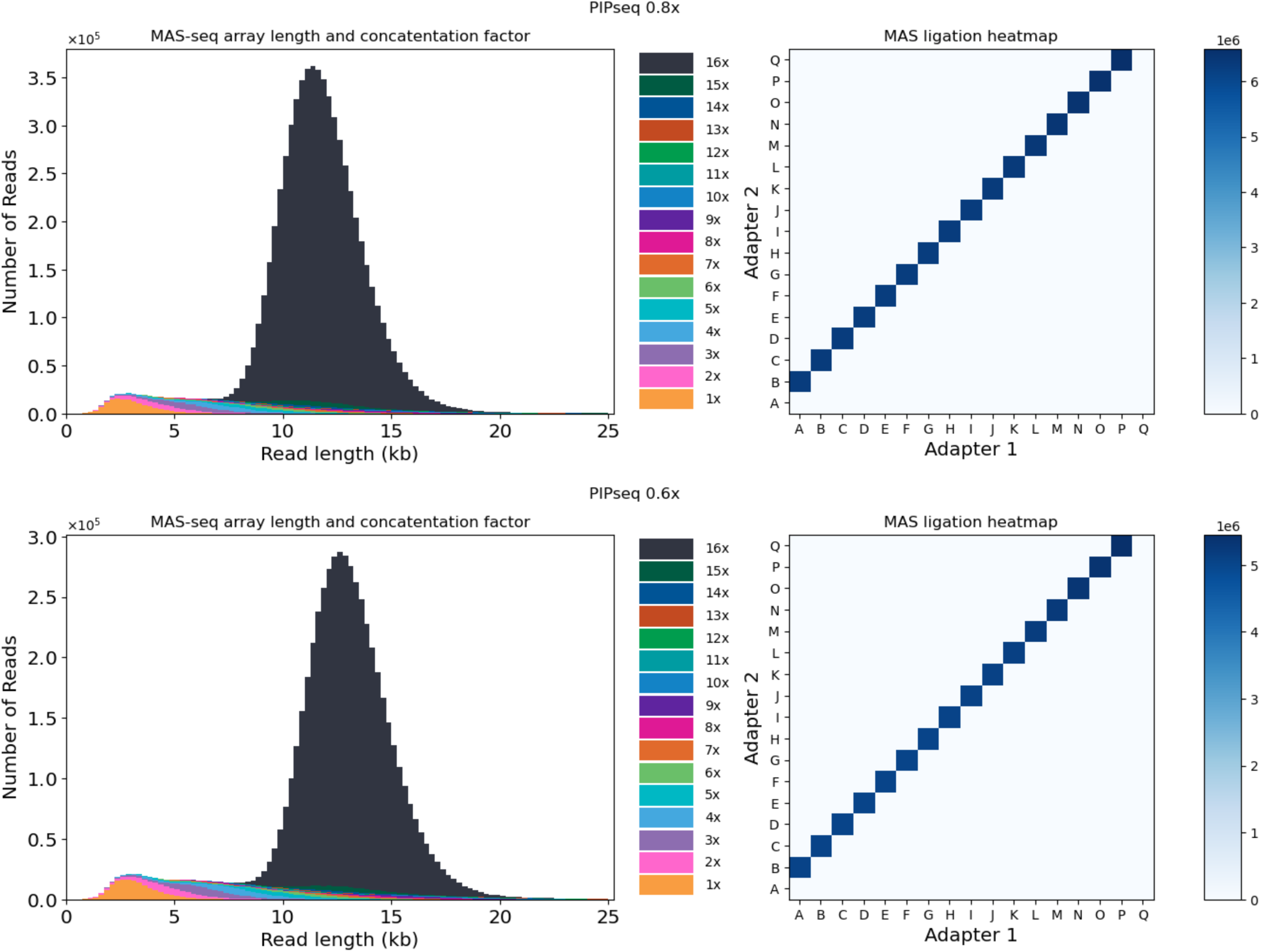
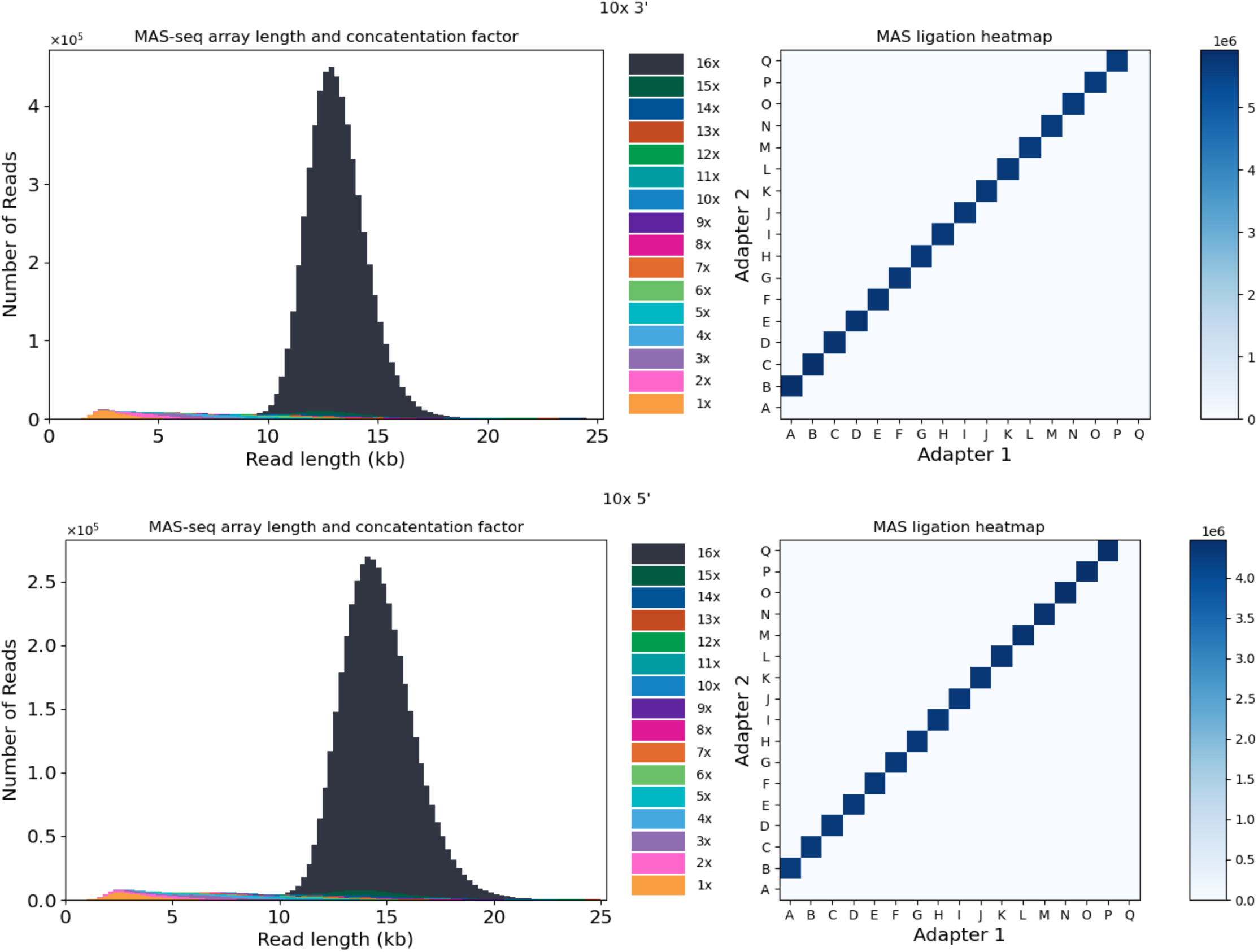
Length histograms and ligation heatmaps for single-cell PBMC Kinnex experiments. 0.8x and 0.6x refer to SPRI ratios used for PIPseq.

**Supplementary Figure 6:**
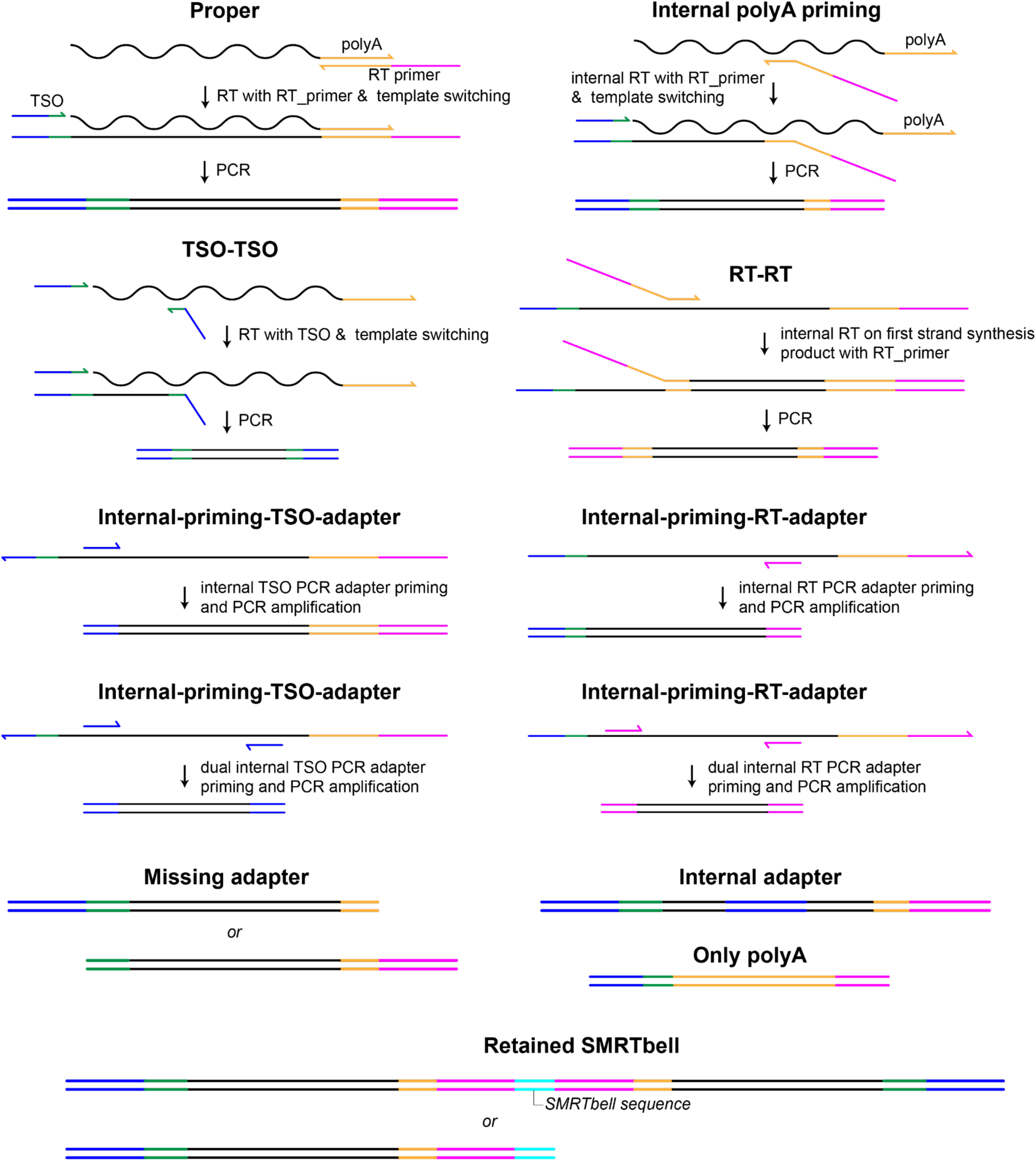
Schematics of artifacts.

**Supplementary Figure 7:**
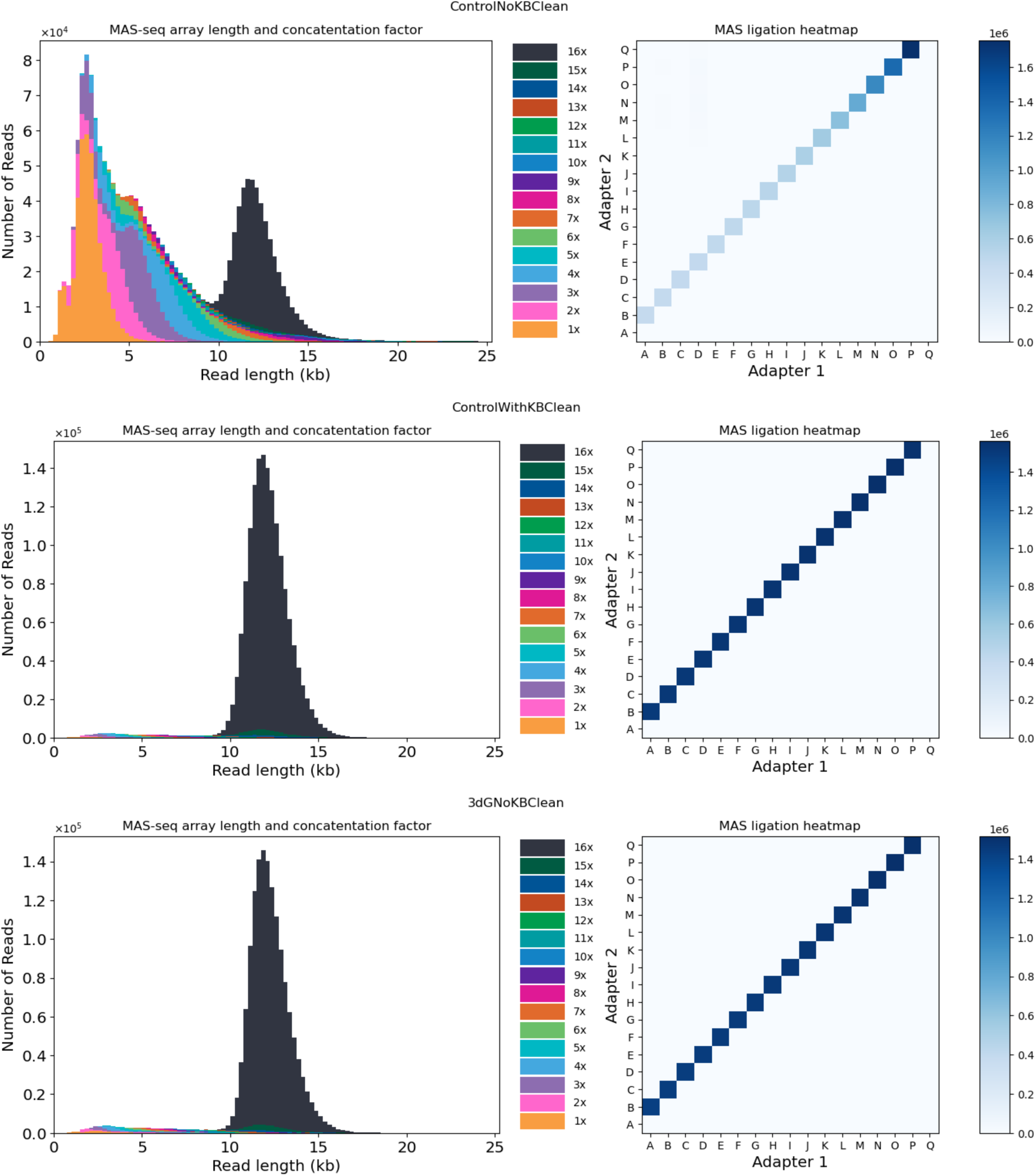
Length histograms and ligation heatmaps for Kinnex data from 10x 3′ (no clean), 10x 3′ (cleaned) and 10x 3dG libraries.

**Supplementary Figure 8:**
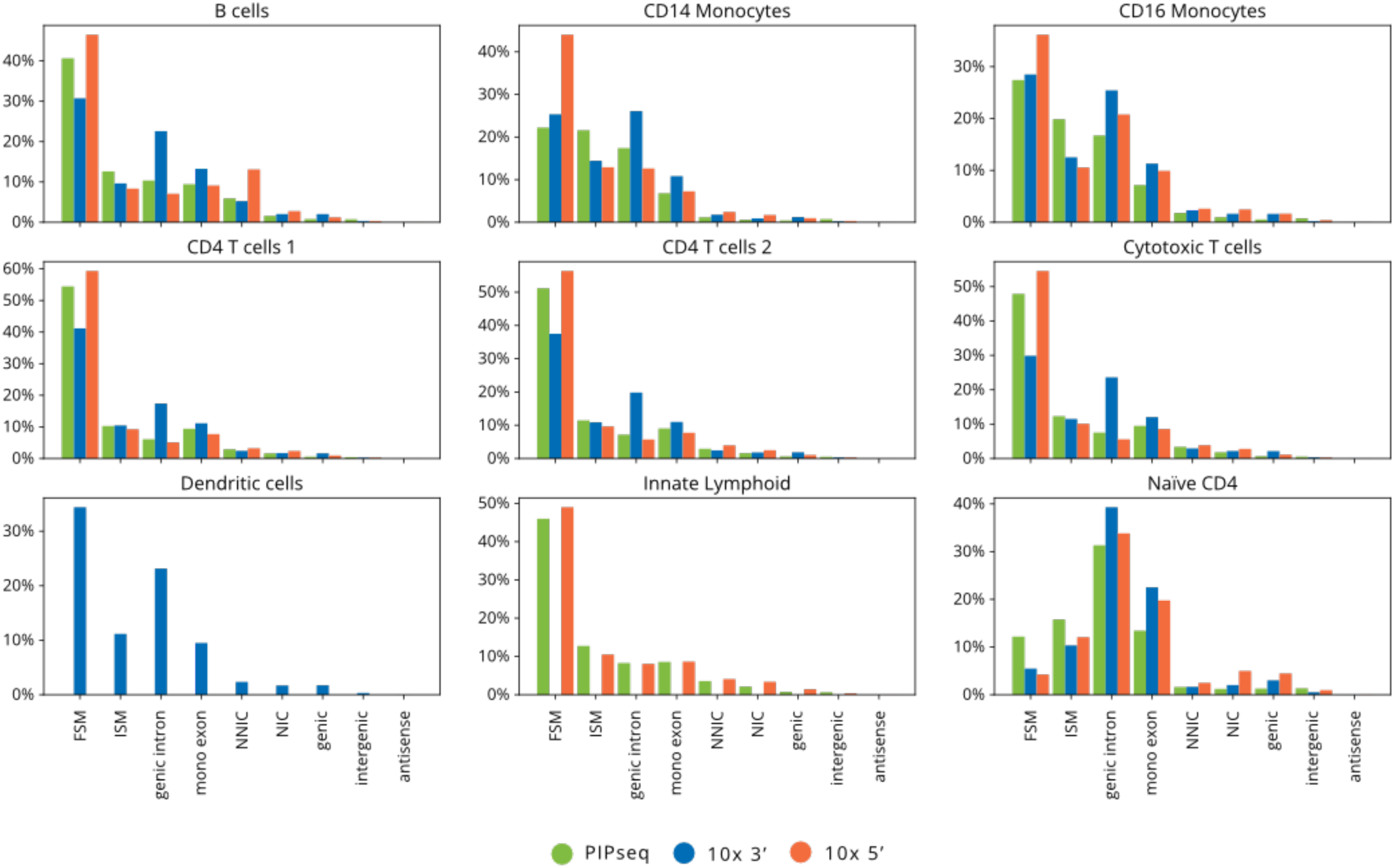
SQANTI3-style classifications for Kinnex data, broken down by cell types determined from the short-read clustering data. FSM: Full splice match; ISM: Incomplete splice match; NNIC: novel not in catalog; NIC: novel in catalog.

**Supplementary Figure 9:**
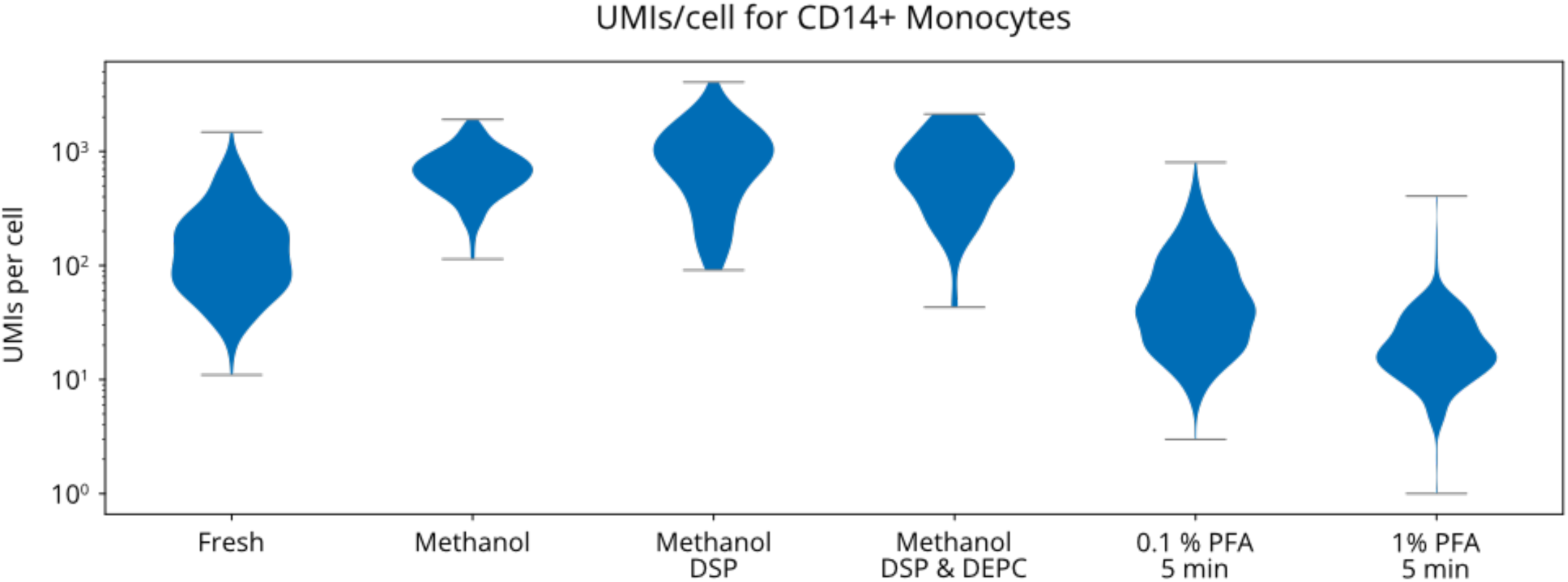
Violin plots showing UMIs/cell for CD14+ monocytes from preliminary comparison of fixation protocols.

**Supplementary Figure 10:**
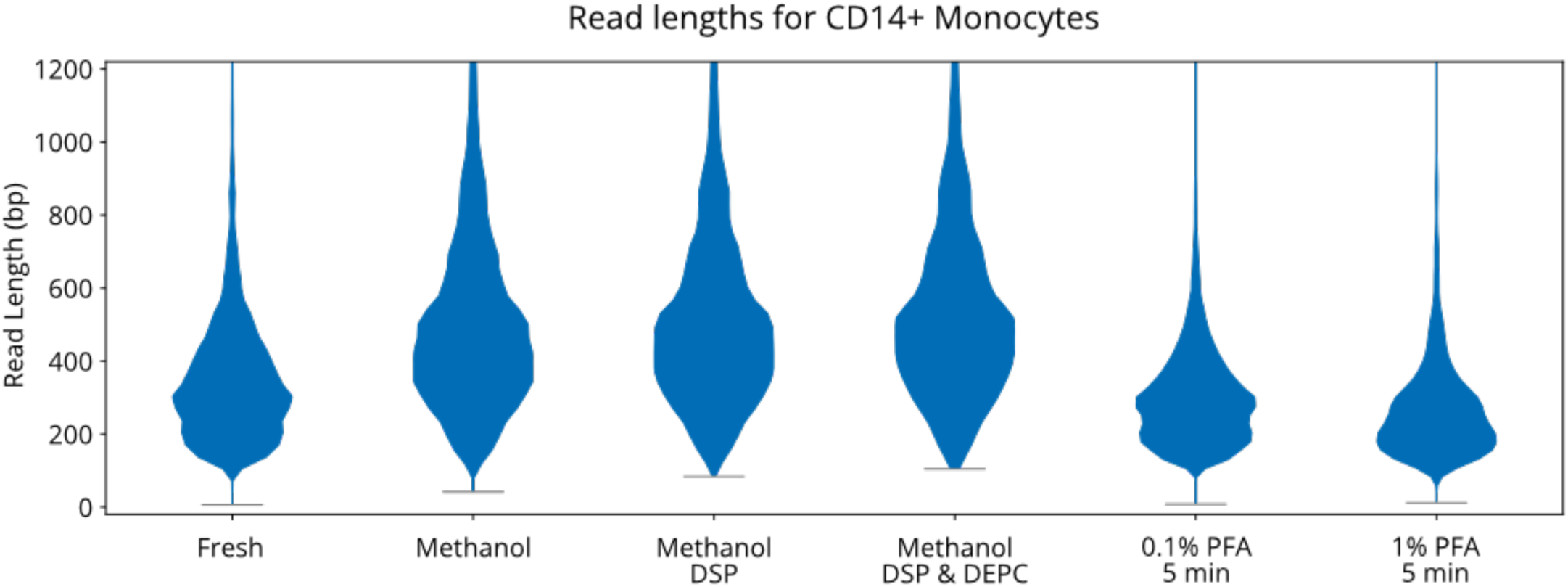
Violin plots showing read length distributions for CD14+ monocytes from preliminary comparison of fixation protocol.

**Supplementary Figure 11:**
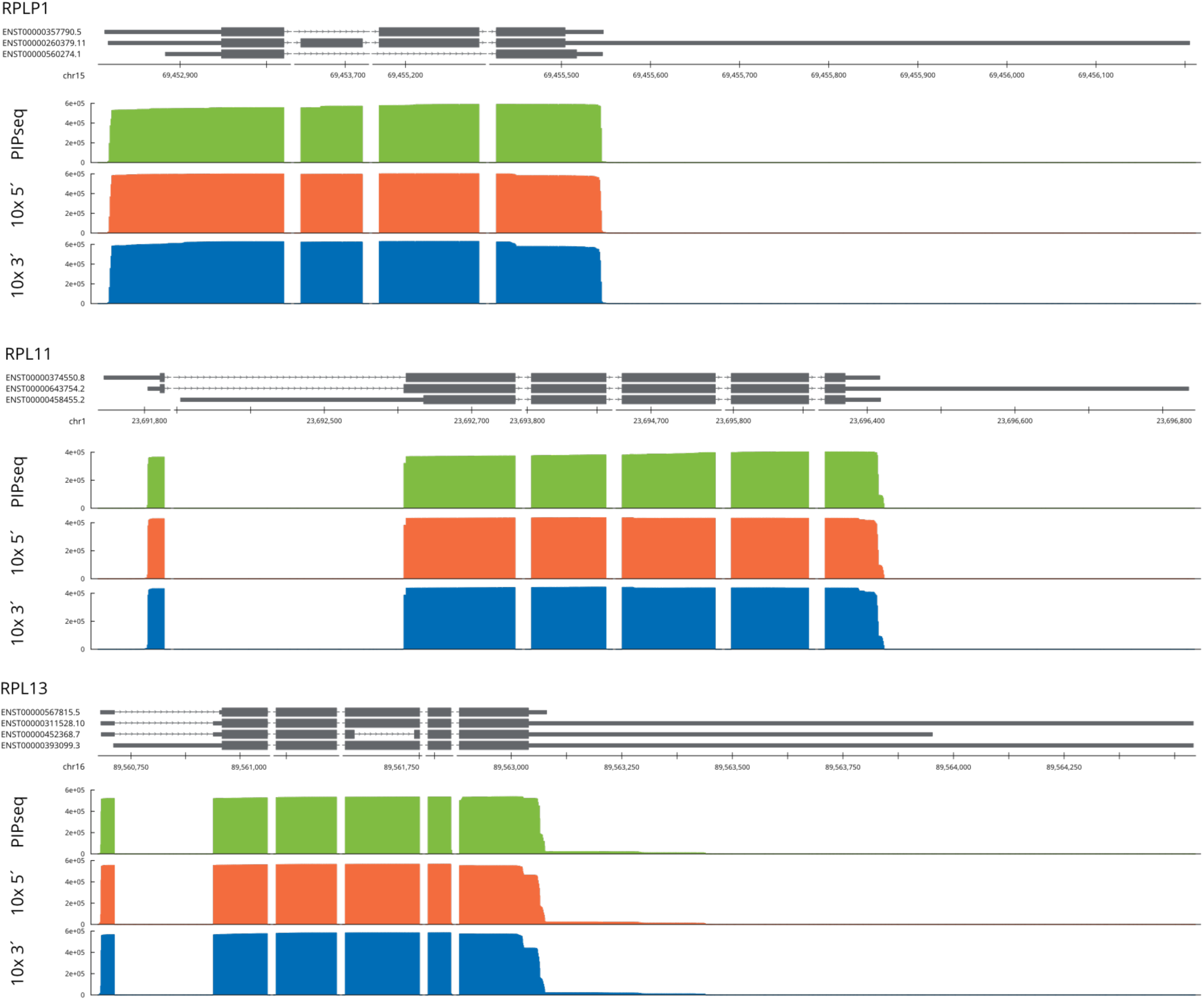
Three examples of alternative polyadenylation sites that lead to transcript misclassification, which contributes to the observed 5′ enrichment in transcript coverage of full splice matches.

**Supplementary Figure 12:**
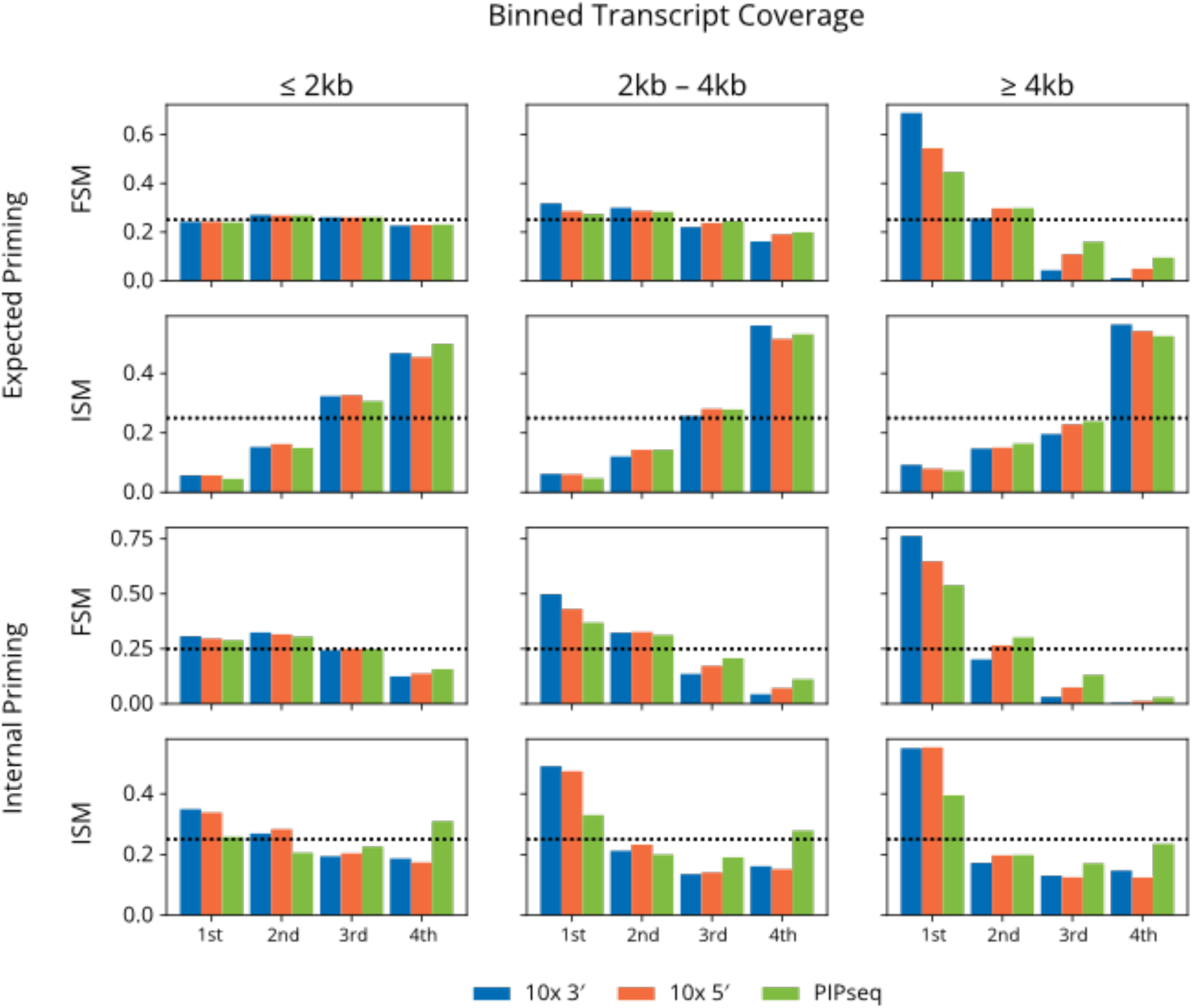
Binned transcript coverage for Kinnex data, broken down by priming classification and splice match. Transcripts have been grouped into three length bins, and average coverage is shown for each quartile of transcript length.

**Supplementary Figure 13:**
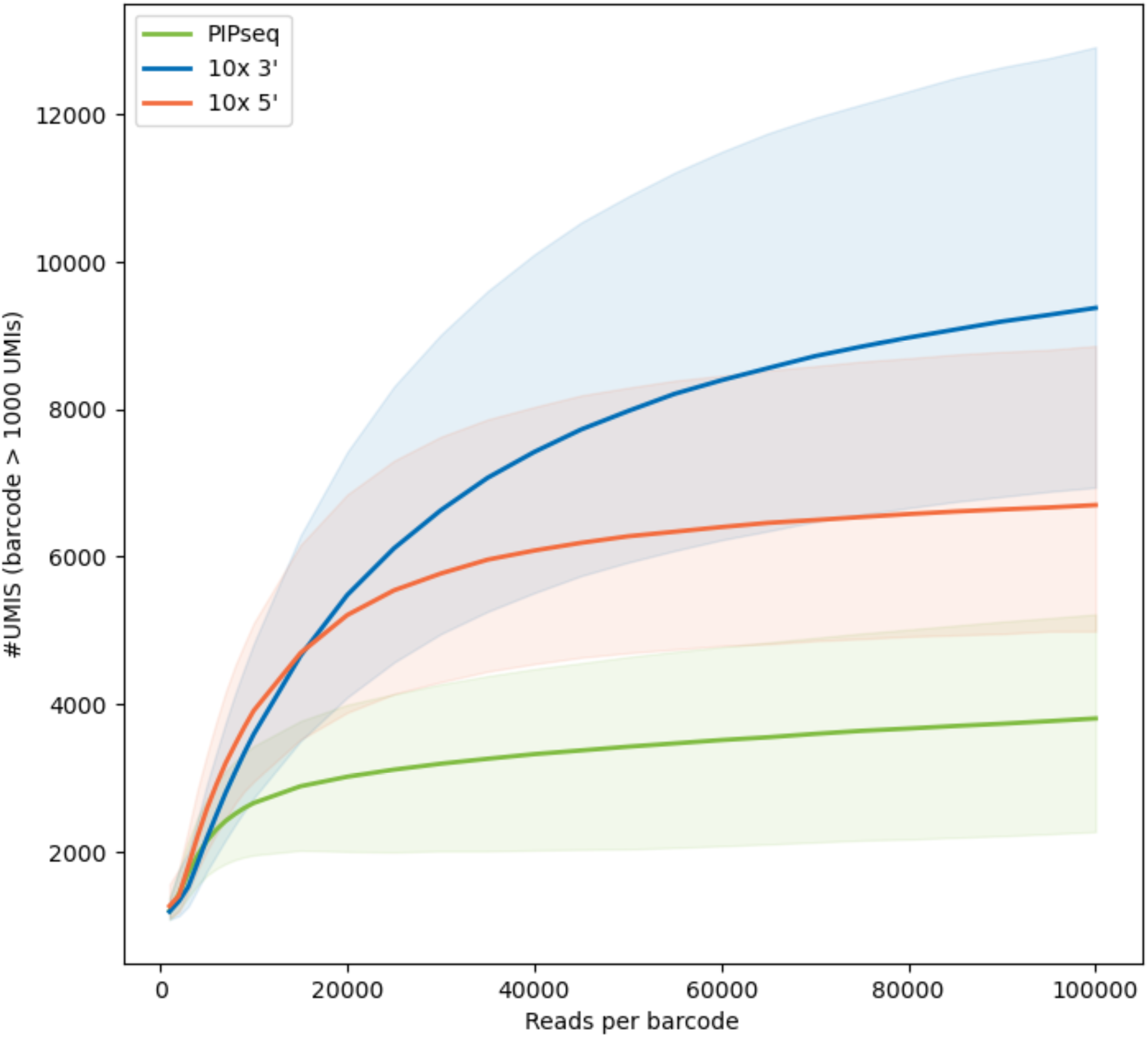
Saturation plot showing the number of UMIs per barcode for barcodes above 1000 UMIs at various subsampling rates.

**Supplementary Figure 14:**
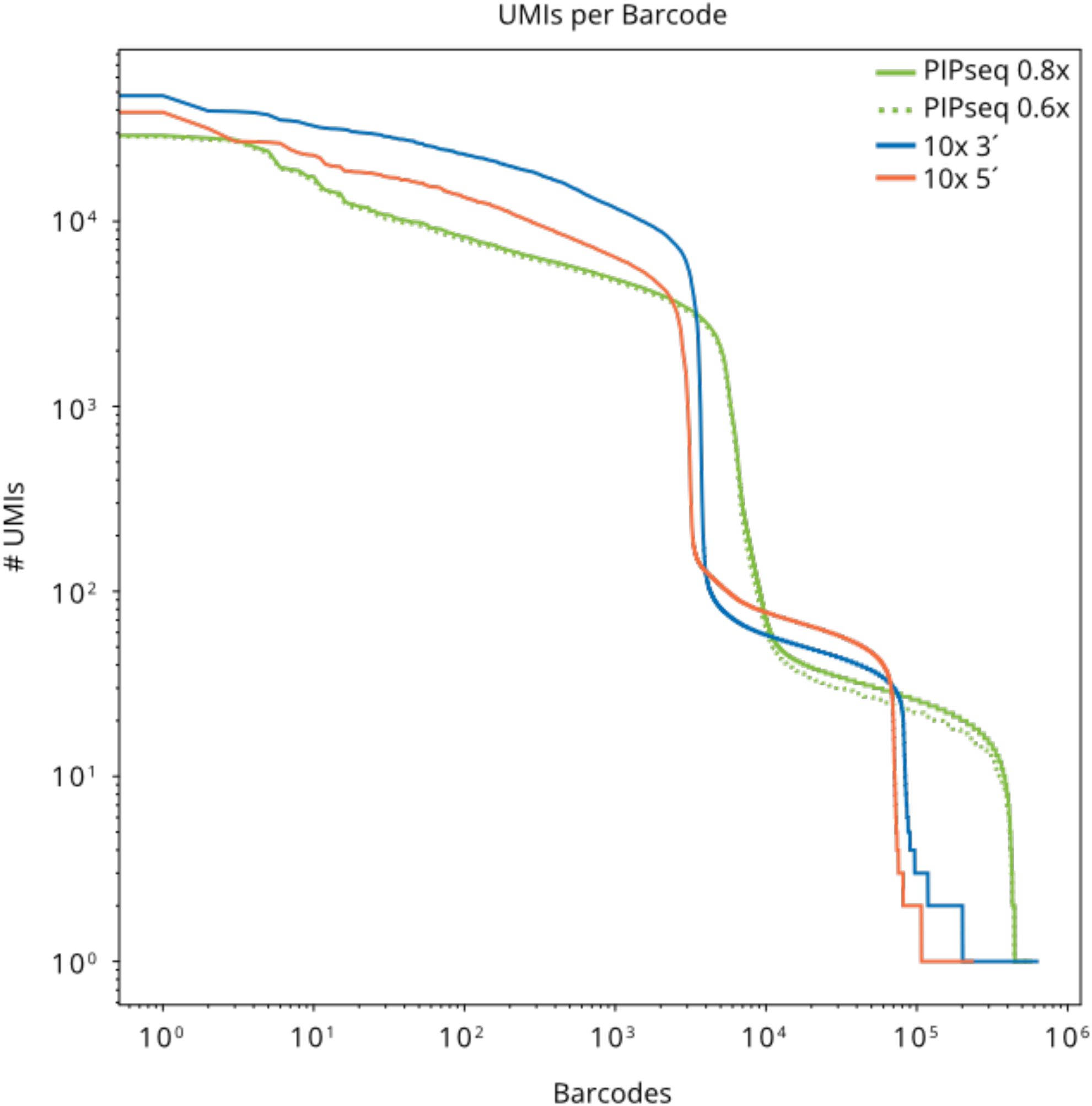
Kneeplots derived from long-read data, similar to Supplementary Fig. 1.

**Supplementary Table 1:** Cell type counts and proportion from short-read clustering.

| Cluster label | PIPseq |  | 10x 3' |  | 10x 5' |  | Notes |
| --- | --- | --- | --- | --- | --- | --- | --- |
|  | % | Count | % | Count | % | Count |  |
| **B cells** | 8.2% | 663 | 7.8% | 289 | 6.7% | 208 | MS4A1+ |
| **CD14 Monocytes** | 8.4% | 682 | 22.3% | 822 | 22.6% | 696 | CD14+ LYZ+ FCGR3A- |
| **CD16 Monocytes** | 18.2% | 1,471 | 6.0% | 222 | 7.0% | 217 | CD14+ LYZ+ FCGR3A+ |
| **CD4 T cells 1** | 25.9% | 2,095 | 29.5% | 1,086 | 30.9% | 952 | CD4+ SELL- |
| **CD4 T cells 2** | 14.7% | 1,190 | 13.3% | 489 | 13.5% | 415 | CD4+ SELL- |
| **Cytotoxic T cells** | 7.9% | 640 | 12.7% | 468 | 7.8% | 241 | CD3+ GNLY+ TRBC1+ |
| **Innate Lymphoid** | 6.1% | 496 | - | - | 7.1% | 220 | NCAM +/- TRDC+ TRBC1+/- |
| **Naive CD4** | 4.9% | 393 | 4.8% | 176 | 4.2% | 131 | CD4+ SELL+ IL7R+ |
| **Dendritic cells** | - | - | 3.4% | 126 | - | - | CLEC9A+ or CD1C+ |
| **Total** |  | **7,630** |  | **3,678** |  | **3,080** |  |

**Supplementary Table 2:**
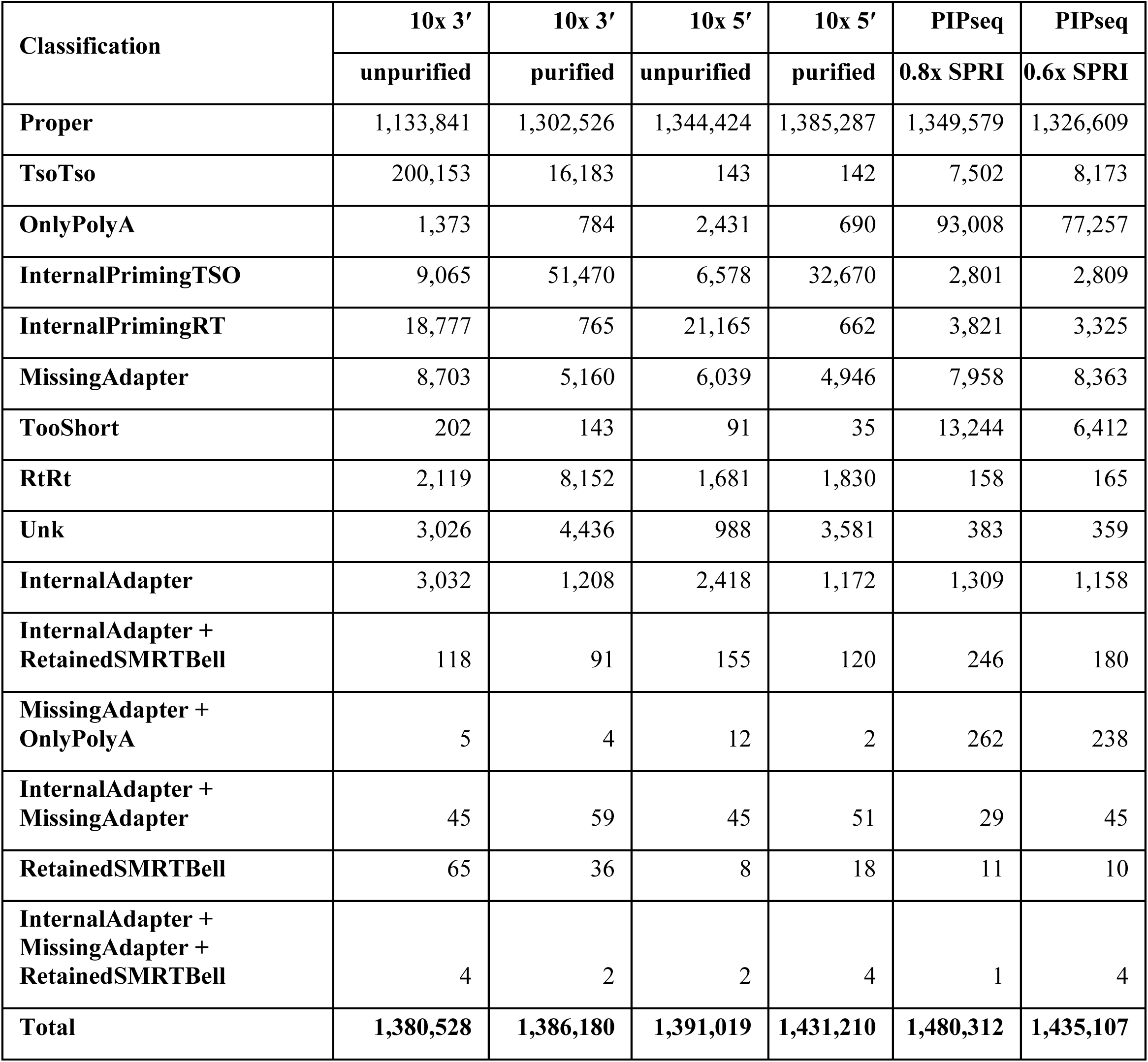
marti classifications from monomer cDNA libraries.

**Supplementary Table 3:** CellRanger metrics for standard TSO and 3dG TSO 10x 3′ experiment.

| Metrics | Control |  | 3dG TSO |  |
| --- | --- | --- | --- | --- |
|  | % | Count | % | Count |
| Estimated Number of Cells |  | 5,417 |  | 5,059 |
| Mean Reads per Cell |  | 31,160 |  | 30,236 |
| Median Genes per Cell |  | 171 |  | 149 |
| Number of Reads |  | 168,794,100 |  | 152,963,325 |
| Valid Barcodes | 98.4% | 166,093,394 | 98.8% | 151,127,765 |
| Sequencing Saturation | 65.7% |  | 74.2% |  |
| Q30 Bases in Barcode | 95.0% |  | 95.0% |  |
| Q30 Bases in RNA Read | 89.4% |  | 89.7% |  |
| Q30 Bases in UMI | 93.5% |  | 93.5% |  |
| Reads Mapped to Genome | 95.1% | 160,523,189 | 95.7% | 146,385,902 |
| Reads Mapped Confidently to Genome | 87.0% | 146,850,867 | 85.0% | 130,018,826 |
| Reads Mapped Confidently to Intergenic Regions | 1.5% | 2,531,912 | 1.0% | 1,529,633 |
| Reads Mapped Confidently to Intronic Regions | 16.1% | 27,175,850 | 7.3% | 11,166,323 |
| Reads Mapped Confidently to Exonic Regions | 69.5% | 117,311,900 | 76.8% | 117,475,834 |
| Reads Mapped Confidently to Transcriptome | 80.0% | 135,035,280 | 81.9% | 125,276,963 |
| Reads Mapped Antisense to Gene | 3.9% | 6,582,970 | 0.7% | 1,070,743 |
| Fraction Reads in Cells | 91.6% | 154,615,396 | 92.1% | 140,879,222 |
| Total Genes Detected |  | 28,687 |  | 24,003 |
| Median UMI Counts per Cell |  | 7,032 |  | 5,072 |

**Supplementary Table 4:**
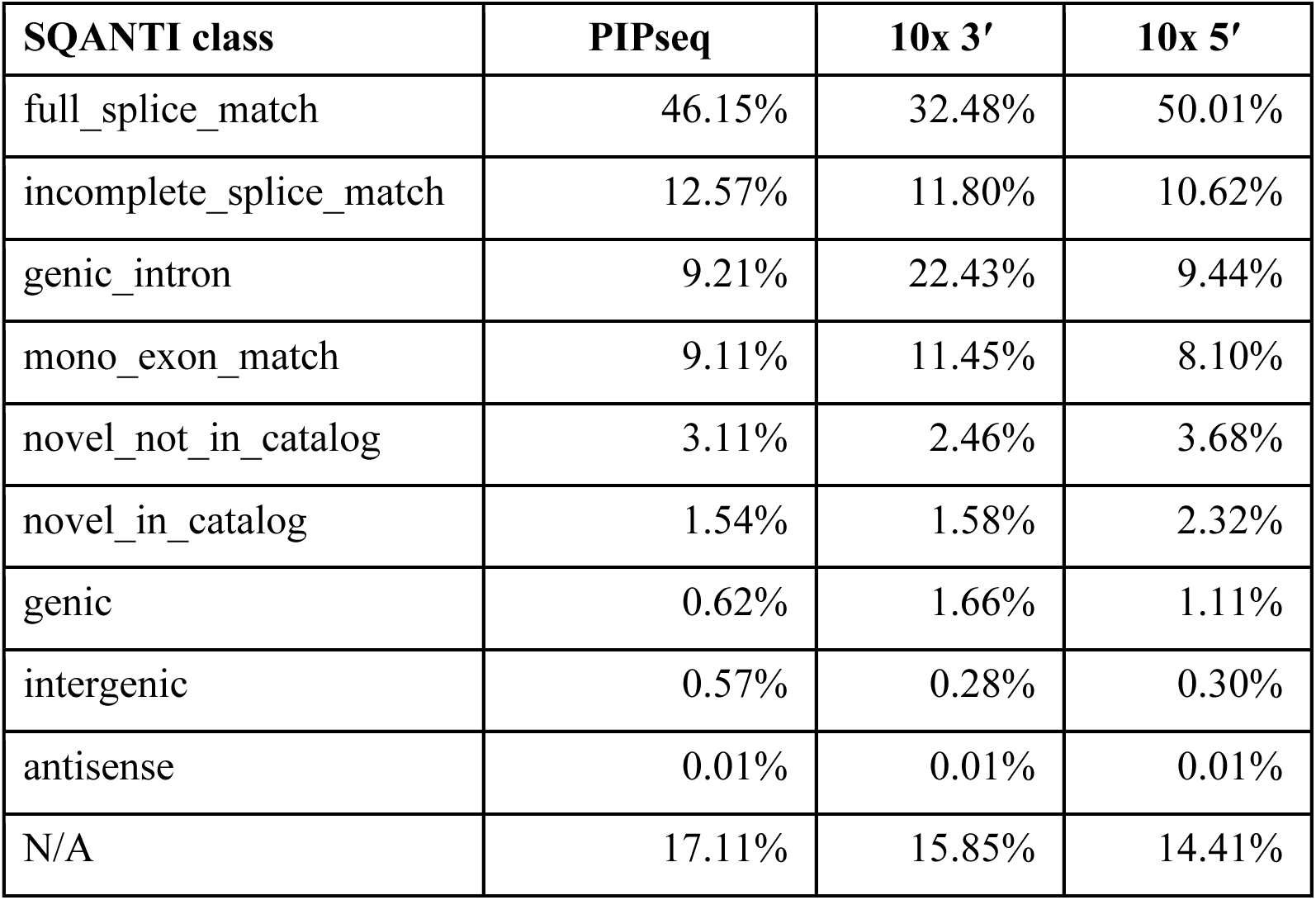
SQANTI classifications from IsoQuant run on primary reads.

| **SQANTI class** | **PIPseq** | **10x 3'** | **10x 5'** |
| --- | --- | --- | --- |
| full_splice_match | 46.15% | 32.48% | 50.01% |
| incomplete_splice_match | 12.57% | 11.80% | 10.62% |
| genic_intron | 9.21% | 22.43% | 9.44% |
| mono_exon_match | 9.11% | 11.45% | 8.10% |
| novel_not_in_catalog | 3.11% | 2.46% | 3.68% |
| novel_in_catalog | 1.54% | 1.58% | 2.32% |
| genic | 0.62% | 1.66% | 1.11% |
| intergenic | 0.57% | 0.28% | 0.30% |
| antisense | 0.01% | 0.01% | 0.01% |
| N/A | 17.11% | 15.85% | 14.41% |

**Supplementary Table 5:** SQANTI classifications from IsoQuant, broken down by cell type.

| **PIPseq** | **CD4 T cells 1** | **CD4 T cells 2** | **Naïve CD4** | **Cytotoxic T cells** | **B cells** | **CD16 Monocytes** | **CD14 Monocytes** | **Innate Lymphoid** |
| --- | --- | --- | --- | --- | --- | --- | --- | --- |
| full_splice_match | 54.45% | 51.21% | 12.58% | 47.87% | 40.65% | 27.48% | 22.01% | 45.97% |
| genic_intron | 6.08% | 7.10% | 30.98% | 7.53% | 10.35% | 16.68% | 17.57% | 8.26% |
| incomplete_splice_match | 10.27% | 11.43% | 15.74% | 12.30% | 12.62% | 19.84% | 21.60% | 12.74% |
| mono_exon_match | 9.40% | 9.03% | 13.34% | 9.47% | 9.42% | 7.16% | 6.79% | 8.56% |
| novel_not_in_catalog | 2.90% | 2.90% | 1.64% | 3.37% | 5.90% | 1.78% | 1.15% | 3.56% |
| novel_in_catalog | 1.60% | 1.56% | 1.22% | 1.80% | 1.57% | 0.96% | 0.57% | 2.10% |
| genic | 0.54% | 0.60% | 1.26% | 0.65% | 0.78% | 0.53% | 0.40% | 0.71% |
| intergenic | 0.44% | 0.50% | 1.31% | 0.53% | 0.73% | 0.73% | 0.73% | 0.62% |
| antisense | 0.01% | 0.01% | 0.02% | 0.01% | 0.01% | 0.01% | 0.01% | 0.01% |
| N/A | 14.31% | 15.66% | 21.91% | 16.47% | 17.98% | 24.83% | 29.18% | 17.47% |

| **10x 3'** | **CD4 T cells 1** | **CD4 T cells 2** | **Naïve CD4** | **Cytotoxic T cells** | **B cells** | **CD14 Monocytes** | **CD16 Monocytes** | **Dendritic cells** |
| --- | --- | --- | --- | --- | --- | --- | --- | --- |
| full_splice_match | 41.18% | 37.51% | 5.47% | 29.90% | 30.74% | 25.33% | 28.49% | 34.40% |
| genic_intron | 17.39% | 19.79% | 39.30% | 23.60% | 22.59% | 26.07% | 25.43% | 23.14% |
| incomplete_splice_match | 10.45% | 10.86% | 10.33% | 11.49% | 9.64% | 14.42% | 12.52% | 11.16% |
| mono_exon_match | 11.13% | 11.00% | 22.49% | 12.02% | 13.26% | 10.82% | 11.30% | 9.49% |
| novel_not_in_catalog | 2.41% | 2.44% | 1.65% | 2.92% | 5.23% | 1.78% | 2.30% | 2.34% |
| novel_in_catalog | 1.71% | 1.81% | 2.02% | 2.17% | 2.03% | 0.91% | 1.63% | 1.69% |
| genic | 1.62% | 1.85% | 3.01% | 2.13% | 2.02% | 1.21% | 1.61% | 1.72% |
| intergenic | 0.26% | 0.31% | 0.51% | 0.35% | 0.33% | 0.24% | 0.24% | 0.32% |
| antisense | 0.01% | 0.01% | 0.02% | 0.01% | 0.01% | 0.01% | 0.01% | 0.01% |
| N/A | 13.83% | 14.41% | 15.20% | 15.42% | 14.15% | 19.21% | 16.48% | 15.73% |

| **10x 5'** | **CD4 T**<br>**cells 1** | **CD4 T**<br>**cells 2** | **Naïve CD4** | **Cytotoxic**<br>**T cells** | **B cells** | **CD14**<br>**Monocytes** | **CD16**<br>**Monocytes** | **Innate**<br>**Lymphoid** |
| --- | --- | --- | --- | --- | --- | --- | --- | --- |
| full_splice_match | 59.22% | 56.58% | 4.22% | 54.44% | 46.48% | 43.99% | 36.14% | 49.07% |
| genic_intron | 5.11% | 5.61% | 33.82% | 5.59% | 7.06% | 12.62% | 20.78% | 8.04% |
| incomplete_splice<br>_match | 9.26% | 9.58% | 12.05% | 10.10% | 8.34% | 12.91% | 10.54% | 10.47% |
| mono_exon_match | 7.72% | 7.71% | 19.77% | 8.58% | 9.10% | 7.26% | 9.91% | 8.60% |
| novel_not_in_catal<br>og | 3.21% | 3.95% | 2.47% | 3.84% | 13.10% | 2.42% | 2.61% | 4.08% |
| novel_in_catalog | 2.37% | 2.48% | 4.97% | 2.79% | 2.75% | 1.70% | 2.47% | 3.37% |
| genic | 0.94% | 1.02% | 4.45% | 1.09% | 1.21% | 0.96% | 1.64% | 1.41% |
| intergenic | 0.25% | 0.27% | 0.95% | 0.27% | 0.32% | 0.31% | 0.40% | 0.32% |
| antisense | 0.01% | 0.01% | 0.03% | 0.01% | 0.01% | 0.01% | 0.01% | 0.01% |
| N/A | 11.91% | 12.80% | 17.25% | 13.29% | 11.61% | 17.83% | 15.50% | 14.63% |

**Supplementary Table 6:**
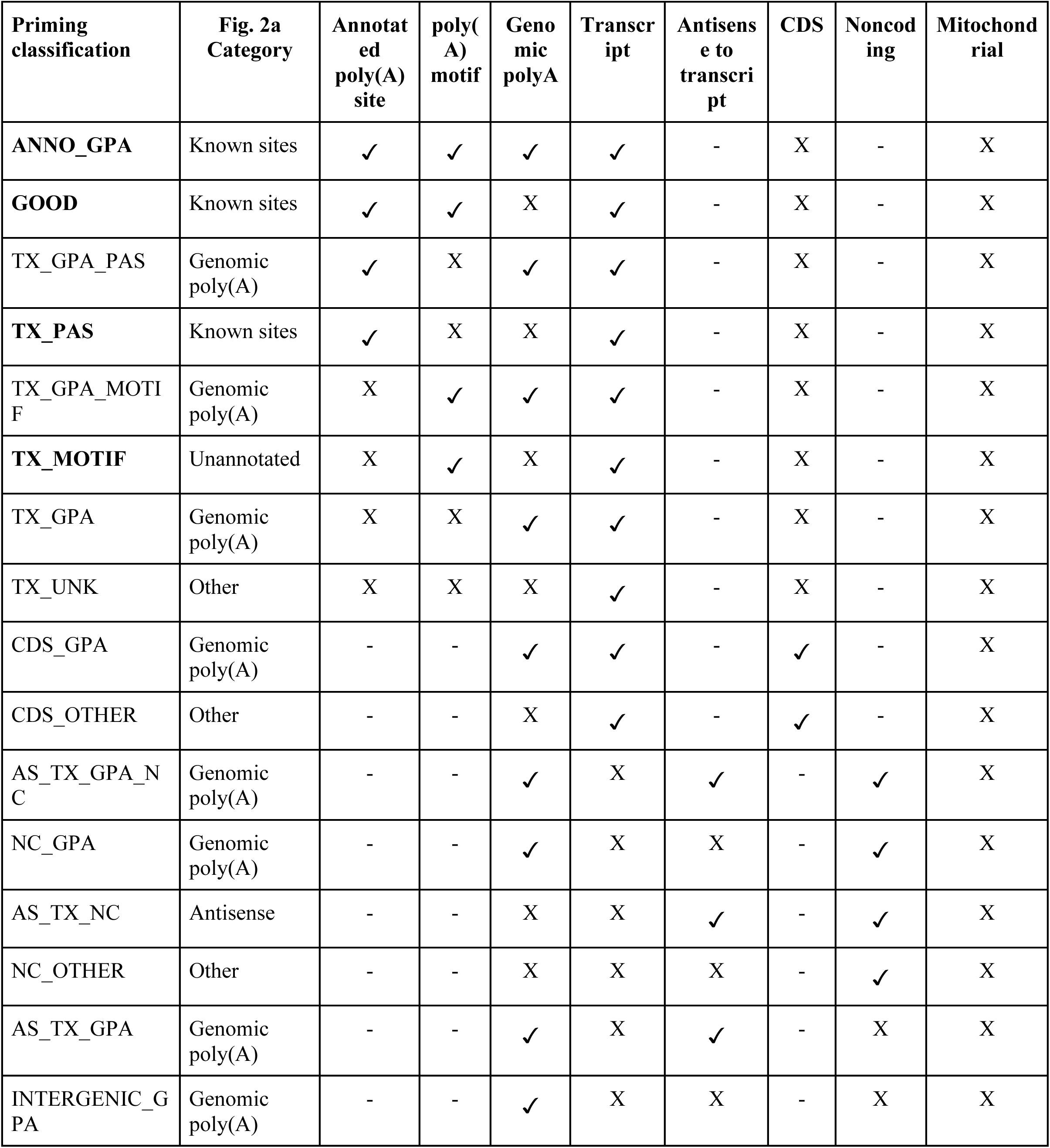
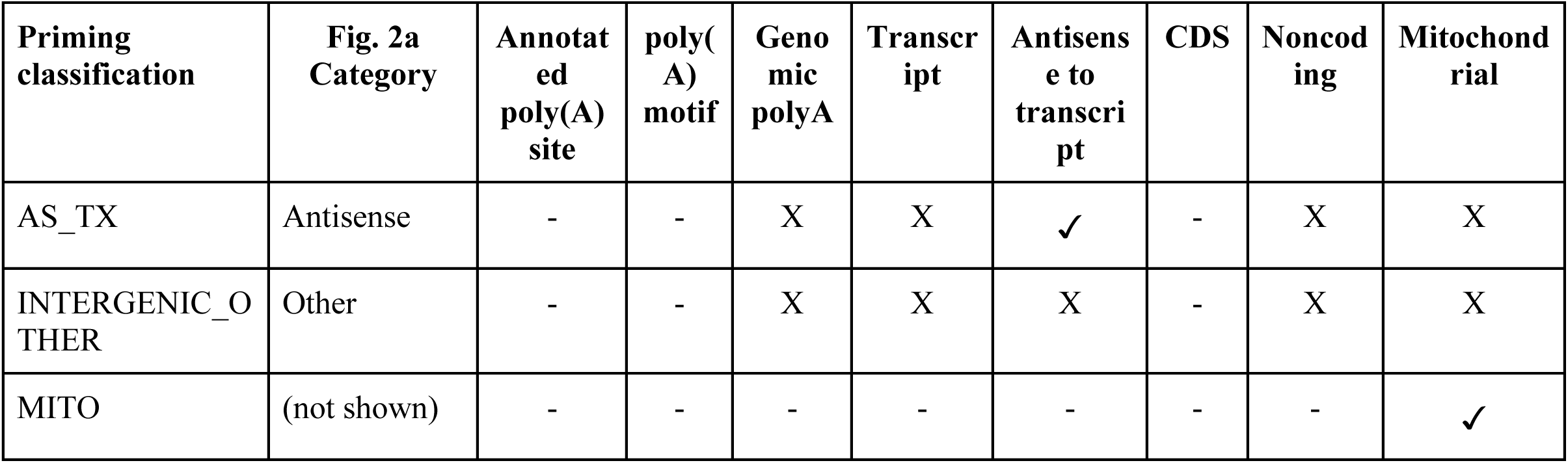
Rules for classification of priming sites based on genomic context of alignment. A check mark indicates a required feature, an X means that feature cannot be present, and a dash means either case is allowed. The categories in bold are expected priming sites, while the rest are considered likely to be mispriming events.

| **Priming classification** | **Fig. 2a Category** | **Annotat ed poly(A) site** | **poly( A) motif** | **Geno mic polyA** | **Transcr ipt** | **Antisens e to transcri pt** | **CDS** | **Noncod ing** | **Mitochond rial** |
| --- | --- | --- | --- | --- | --- | --- | --- | --- | --- |
| **ANNO_GPA** | Known sites | ✓ | ✓ | ✓ | ✓ | - | X | - | X |
| **GOOD** | Known sites | ✓ | ✓ | X | ✓ | - | X | - | X |
| TX_GPA_PAS | Genomic poly(A) | ✓ | X | ✓ | ✓ | - | X | - | X |
| **TX_PAS** | Known sites | ✓ | X | X | ✓ | - | X | - | X |
| TX_GPA_MOTIF | Genomic poly(A) | X | ✓ | ✓ | ✓ | - | X | - | X |
| **TX_MOTIF** | Unannotated | X | ✓ | X | ✓ | - | X | - | X |
| TX_GPA | Genomic poly(A) | X | X | ✓ | ✓ | - | X | - | X |
| TX_UNK | Other | X | X | X | ✓ | - | X | - | X |
| CDS_GPA | Genomic poly(A) | - | - | ✓ | ✓ | - | ✓ | - | X |
| CDS_OTHER | Other | - | - | X | ✓ | - | ✓ | - | X |
| AS_TX_GPA_NC | Genomic poly(A) | - | - | ✓ | X | ✓ | - | ✓ | X |
| NC_GPA | Genomic poly(A) | - | - | ✓ | X | X | - | ✓ | X |
| AS_TX_NC | Antisense | - | - | X | X | ✓ | - | ✓ | X |
| NC_OTHER | Other | - | - | X | X | X | - | ✓ | X |
| AS_TX_GPA | Genomic poly(A) | - | - | ✓ | X | ✓ | - | X | X |
| INTERGENIC_GPA | Genomic poly(A) | - | - | ✓ | X | X | - | X | X |

| Priming classification | Fig. 2a Category | Annotat ed poly(A) site | poly( A) motif | Geno mic polyA | Transcr ipt | Antisens e to transcri pt | CDS | Noncod ing | Mitochond rial |
| --- | --- | --- | --- | --- | --- | --- | --- | --- | --- |
| AS_TX | Antisense | - | - | X | X | ✓ | - | X | X |
| INTERGENIC_OTHER | Other | - | - | X | X | X | - | X | X |
| MITO | (not shown) | - | - | - | - | - | - | - | ✓ |

**Supplementary Table 7:** Full priming classification, not including mitochondrial reads.

| **Priming classification** | **PIPseq** | **10x 3'** | **10x 5'** |
| --- | --- | --- | --- |
| **ANNO_GPA** | 0.76% | 0.53% | 0.55% |
| **GOOD** | 60.48% | 40.21% | 52.75% |
| TX_GPA_PAS | 0.01% | 0.03% | 0.03% |
| **TX_PAS** | 1.55% | 0.99% | 1.40% |
| TX_GPA_MOTIF | 7.02% | 10.63% | 8.76% |
| **TX_MOTIF** | 11.15% | 11.27% | 9.88% |
| TX_GPA | 3.38% | 2.97% | 2.35% |
| TX_UNK | 1.30% | 4.12% | 2.39% |
| CDS_GPA | 1.23% | 2.03% | 2.34% |
| CDS_OTHER | 1.06% | 4.86% | 4.52% |
| AS_TX_GPA_NC | 0.13% | 0.35% | 0.14% |
| NC_GPA | 1.73% | 1.99% | 1.74% |
| AS_TX_NC | 0.27% | 0.31% | 0.19% |
| NC_OTHER | 2.56% | 6.10% | 6.98% |
| AS_TX_GPA | 1.27% | 6.05% | 1.54% |
| INTERGENIC_GPA | 3.50% | 2.71% | 1.79% |
| AS_TX | 0.38% | 2.65% | 0.46% |
| INTERGENIC_OTHER | 2.24% | 2.23% | 2.20% |
| **Total** | **64,051,269** | **70,709,925** | **53,602,563** |

**Supplementary Table 8:**
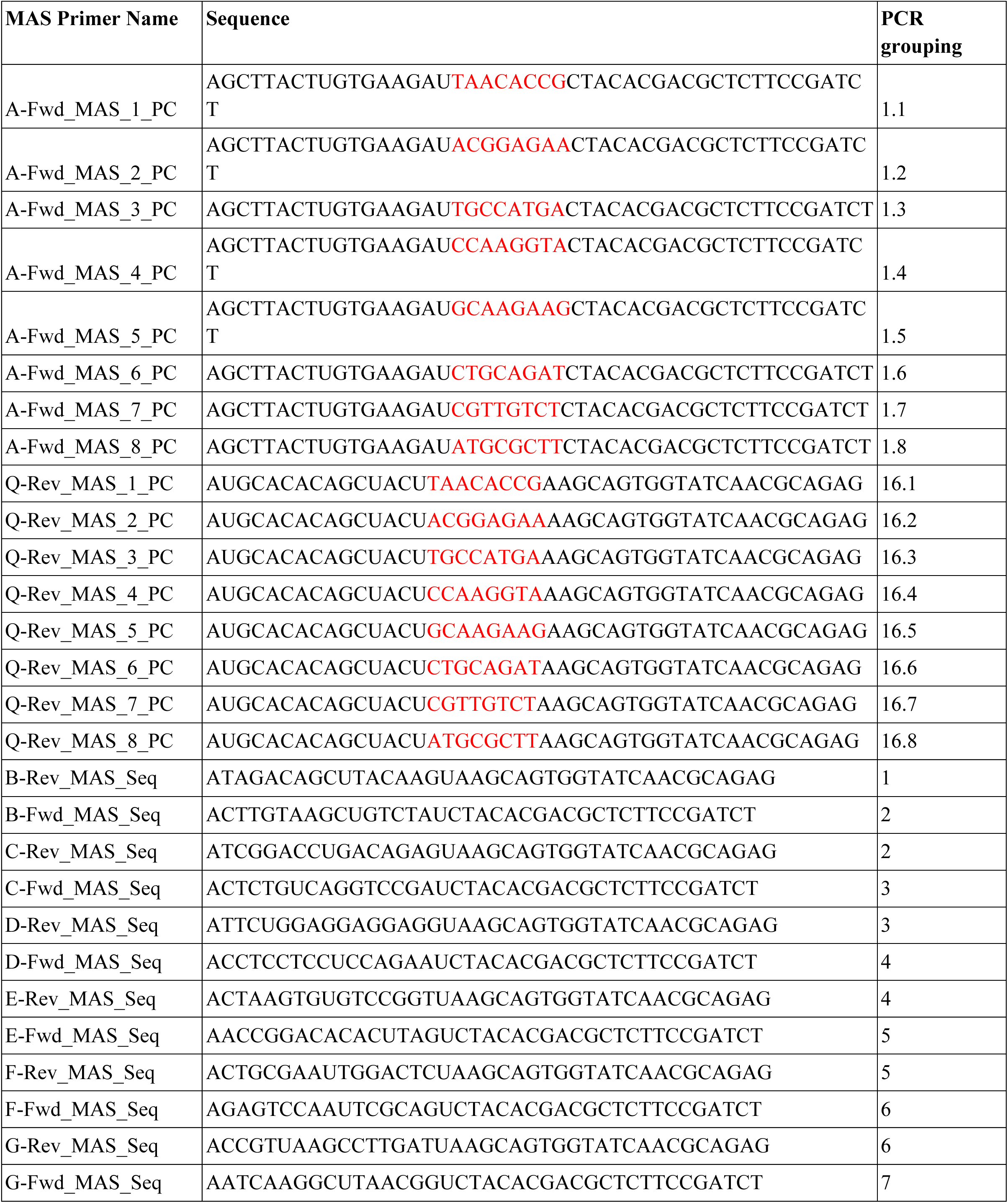
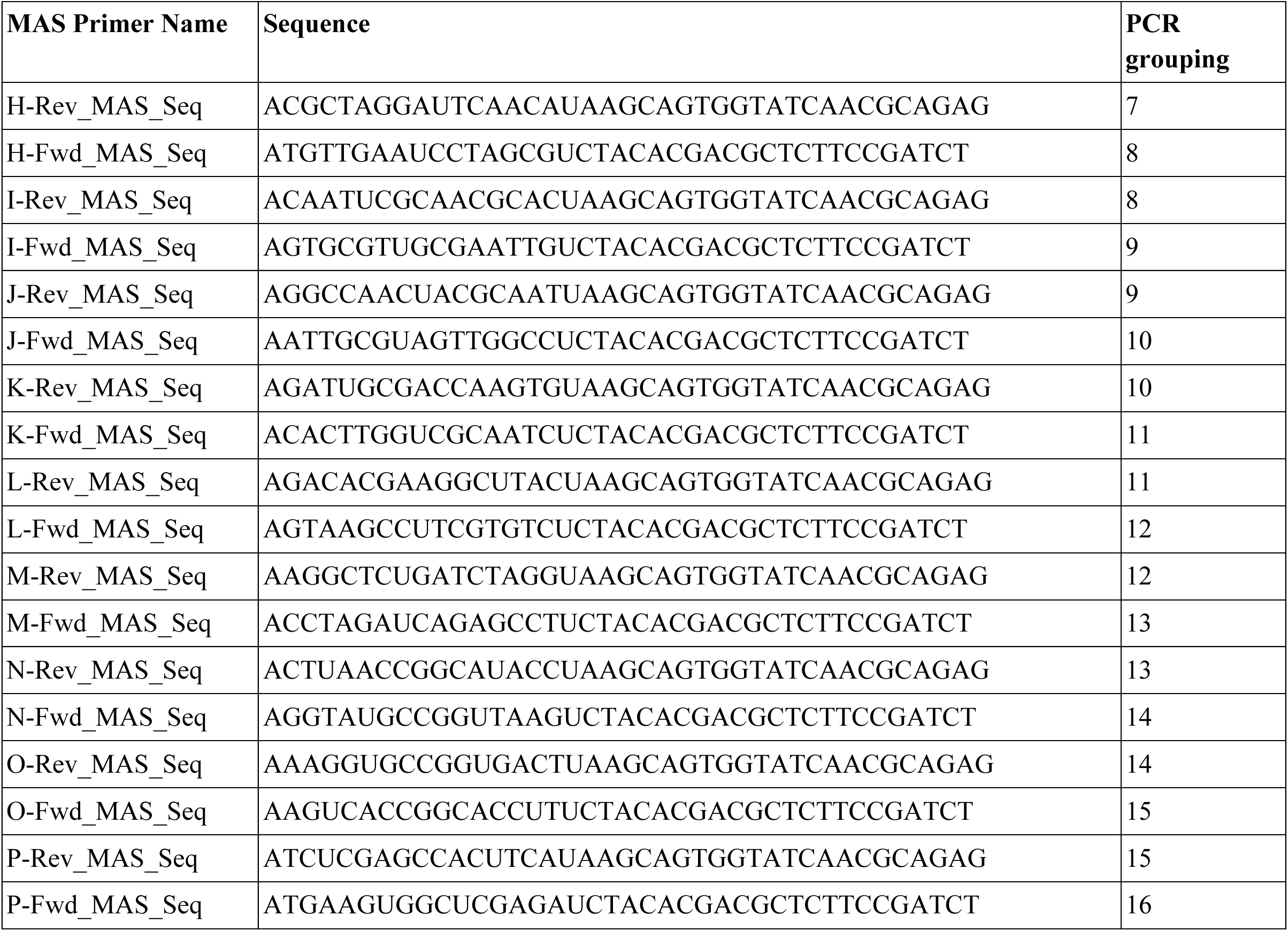
MAS Primer Sequences.

